# Benzothiazole Derivatives as Dual Modulators of PGE2 and GABAergic Signaling in Skeletal Muscle

**DOI:** 10.64898/2026.05.30.728982

**Authors:** Marian N. Aziz, Kamal Awad, Jian Huang, Zhiying Wang, Venu Varanasi, Marco Brotto, Carl J. Lovely

## Abstract

Benzothiazoles are attractive scaffolds for small-molecule modulators of neuronal signaling. However, their impact on skeletal muscle and GABAergic pathways remains poorly understood. We synthesized a focused library of benzothiazole derivatives via oxidative electrophilic substitution and profiled their activity in C2C12 skeletal muscle cells, assessing cytotoxicity, proliferation, myogenic differentiation, and GABA-related signaling using cell-based assays, real-time PCR, and transcriptomics. Omics-guided analyses revealed that selected benzothiazole derivatives differentially modulate myogenic differentiation and prostaglandin E2, and simultaneously bidirectionally regulate GABAergic and glutamatergic signaling genes, including synaptic subunits and transporters. Notably, a lead derivative downregulated Gabrg2, a GABA-A receptor subunit implicated in epilepsy and other disorders of inhibitory synapses, highlighting a potential link between skeletal muscle signaling and neuropsychiatric disease. These findings position benzothiazole derivatives as candidate modulators of GABAergic signaling with translational potential for conditions involving dysfunctional inhibitory synapses.

## 1. INTRODUCTION

Gamma-Aminobutyric acid (GABA) is an amino acid neurotransmitter synthesized by glutamic acid decarboxylase enzyme via the decarboxylation of glutamate. Since 1958, GABA has been classified as a significant neurotransmitter inhibitor, and GABA aminotransferase was identified as a therapeutic target for several neurological diseases.(*1–3*) This small natural activator, GABA, mediates synaptic inhibition via binding to GABA receptors (A and B), which function as receptor-gated chloride channels.(*4*) GABA-regulated chloride gates allow chloride ions to pass through neurons, resulting in hyperpolarization of these negative charges, thus reducing the probability of changes in neuron action potentials.(*5*) GABA-A receptor is an ionotropic and heteropentameric channel consisting of five subunits (two α_1_, two β_2_, one γ_2_). This subunit composition represents the most common GABA isoform in the mammalian central nervous system (CNS) and is encoded by Gabra1, Gabrb2, and Gabrg2 genes.(*6–8*) Attention has mainly been focused on the GABA-A receptor due to its at least three allosteric binding sites, making it a pharmacologically complex receptor for treating various central nervous system diseases. These include the benzodiazepine site for reducing epileptic seizures, the sedative site for anxiety treatment, and the site of the depressant barbiturates.(*8*) The γ_2_ subunits of the GABA type A gene play a major role in the maintenance of GABA-A receptors at mature synapses, and their deletion resulted in the absence of a benzodiazepine binding site.(*9*) Over half of the known epilepsy-associated mutations have been identified in GABAG2.(*10, 11*) Therefore, the Gabrg2 gene is considered an epilepsy gene and is a potential molecular target for the discovery of novel therapeutic strategies for genetic and non-genetic epilepsy and other diseases associated with a mutation in GABAergic synapses.(*12*) Furthermore, GABAergic system dysfunction is associated with several muscle-related disorders, including hyperekplexia, GABA-transaminase deficiency, stiff-person syndrome, tetanus, strychnine poisoning, and muscle symptoms associated with Huntington’s and Parkinson’s diseases.(*13–16*) On the other hand, solute carrier family 7 member 11 (SLC7α11, system Xc-, glutamate transporter) is one of the important plasma membrane transporter systems in humans, regulating the intracellular glutathione level.(*17, 18*) The SLC7α11 gene encodes a Na-independent antiporter/exchanger that reduces extracellular L-cystine (a dimer of cysteine) into cysteine, a key intermediate in intracellular L-glutathione synthesis.(*19*) Thus, it protects the cells from reactive oxygen species (ROS) and converts toxic lipids into healthy lipids.(*17*) Furthermore, the neurotransmitter glutamate plays a significant role in the functioning of the central nervous system, and its dysfunction is associated with several CNS disorders.(*20–22*)

A growing body of research has also established prostaglandin E2 (PGE_2_), a bioactive lipid mediator generated by the cyclooxygenase (COX) pathway, as a critical regulator of skeletal muscle myogenesis, regeneration, and repair.(*23–26*) PGE_2_ exerts direct effects on muscle stem cells (MuSCs) and myoblasts, primarily via the EP4 receptor, where it promotes proliferation, differentiation, mediates inflammatory signaling following muscle injury, and modulates oxidative stress.(*24–28*) Altered levels of PGE_2_ in the muscle microenvironment can enhance or impair muscle regeneration, influence muscle strength, and affect myogenic progression.(*26–28*) The importance of PGE_2_ as a regulator for MuSC expansion and muscle repair underlies an emerging connection between lipid mediators and myogenic regulatory programs.

By classical definition, benzothiazoles are a class of heterocyclic molecules that contain a substituted phenyl ring fused with a five-member ring containing sulfur and nitrogen atoms, called a thiazole moiety.(*29*) Benzothiazoles are core structures in many bioactive and drug molecules with different pharmacological activities. **Figure 1** presents the approved drugs that contain benzothiazole (thiazole) derivatives. Riluzole (**1**) is the most commonly known benzothiazole categorized under CNS medicines for the treatment of amyotrophic lateral sclerosis (ALS).(*30*) Thus, numerous benzothiazole-based analogs have been developed in attempts to improve their corresponding activity against ALS.(*30*) In addition, the benzothiazole motif serves as the core structure in many approved drugs, such as probenzole (**2**) (agrochemical drug),(*31*) ethoxazolamide (**3**) (carbonic anhydrase inhibitor for glaucoma treatment),(*32*) and zopolrestat (**4**) (aldose reductase inhibitor for treating diabetic complications).(*33*) Additionally, bioactivity can be driven by the thiazole ring, which is found in numerous approved drugs, such as pramipexole (**5**), a tetrahydrobenzothiazole, which can be used to treat Parkinson’s disease and restless leg syndrome.(*34, 35*) Thus, there is great interest in developing and investigating biologically active benzothiazole derivatives. As a result, numerous studies and patents have described various derivatives based on the benzothiazole scaffold.(*29*) To name a few, benzothiazoles serve as anticonvulsants,(*36–40*) anti-inflammatories,(*41–43*) anti-proliferative agents,(*44–48*) antimicrobials,(*49, 50*) anti-analgesics(*51–53*), and antioxidant agents.(*54–56*) Due to the broad potential bioactivity of benzothiazoles, organic and medicinal chemists are interested in discovering alternative synthetic methodologies and investigating the biological activity of novel synthesized benzothiazole derivatives.

**Figure 1.**
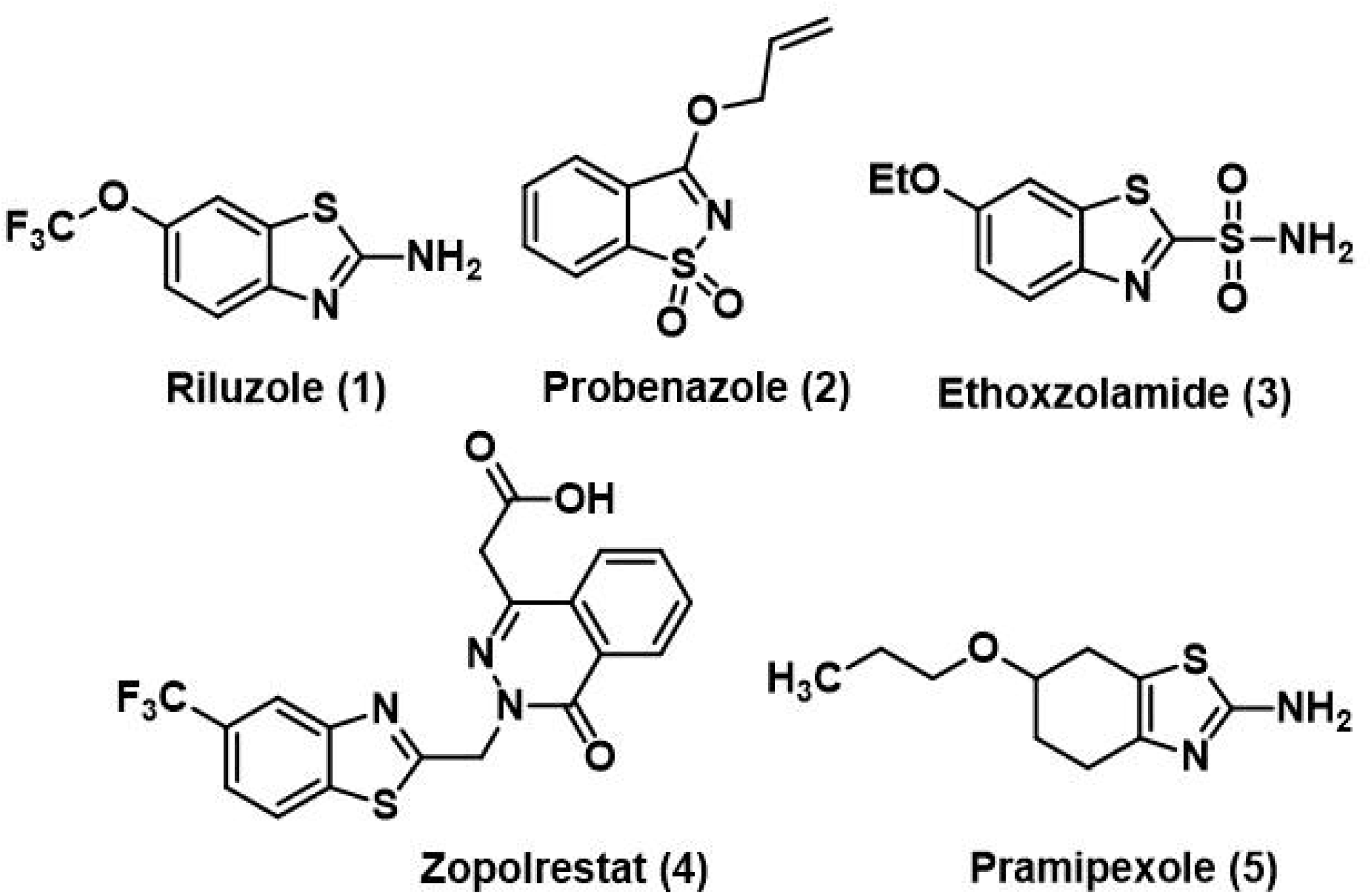
Approved drugs contain benzothiazole (thiazole) derivatives (1–5).

The oxidative S-cyclization of N-aryl thiourea derivatives yields 2-aminobenzothiazoles, which is a convenient method for constructing thiazoles. Many approaches have been studied for this direct transformation using metal-catalyzed oxidative cyclization, such as ruthenium chloride,(*57*) palladium acetate,(*58*) and nickel salts.(*59*) 2-Aminobenzothiazoles can also be synthesized using aniline derivatives with different types of amines and thiocarbonyl sources via various oxidative conditions. These conditions include iodine- or bromine-induced oxidations,^48^ Copper-catalyzed Ullmann-type reactions,(*60*) thiocarbamoyl as a source for the thioamide group,(*58, 61, 62*) and three-component reactions using potassium sulfide mediated by copper chloride.(*63*)

In the present study, a new synthetic methodology for producing novel functionalized benzothiazoles is described, together with an exploration of their effects on muscle myogenesis, GABAergic signaling, and, for the first time, the mechanistic regulation of the GABA-PGE_2_ axis, highlighting new biological opportunities at the intersection of neurotransmitter and prostaglandin signaling in muscle for therapeutic innovation.

## 2. MATERIALS AND METHODS

### 2.1 Experimental Design

The primary objective of this study was to synthesize and characterize a series of thiourea and benzothiazole derivatives and to evaluate their effects on skeletal muscle cell viability, differentiation, redox status, and metabolomic and lipidomic profiles. Thiourea derivatives were prepared by reacting 1-(4-methoxyphenyl)-N-methylmethanamine with aryl isothiocyanates under mild conditions, followed by oxidative cyclization using iodosobenzene diacetate (IBDA) to yield benzothiazole derivatives. All intermediates and final compounds were purified using flash chromatography and characterized by NMR, IR spectroscopy, and HR-MS spectrometry.

For biological evaluation, murine C2C12 skeletal myoblasts were employed as an in vitro model. Cells were cultured under standard conditions and exposed to benzothiazole derivatives at varying concentrations (5–100 µM) to assess cytotoxicity using the MTS assay. Differentiation studies were conducted by culturing cells in differentiation medium supplemented with 5 µM or 10 µM of selected compounds (**7b** and **7g**). Morphological differentiation was assessed via immunostaining for myosin heavy chain (MHC) and nuclear counterstaining with DAPI.

Metabolic profiling included quantification of γ-aminobutyric acid (GABA) and related amino acid metabolites (AABA, BABA, BAIBA) in both cells and conditioned media using a validated LC–MS/MS method. Additionally, global lipid mediator profiling was performed using an LC–ESI–MS/MS-based lipidomics approach optimized for 158 lipid mediators. Gene expression analyses were performed on RNA isolated from differentiated C2C12 cells treated with 10 µM of **7b** or **7g** using the Mouse GABA & Glutamate RT² Profiler PCR Array. Oxidative stress was assessed by fluorescence imaging after carboxy-H₂DCFDA staining, and relative ROS intensity was quantified using ImageJ software. Differential gene and metabolite data were further interpreted through Ingenuity Pathway Analysis (IPA) to identify affected signaling pathways and molecular networks.

### 2.2 Chemical synthesis

All reagents were purchased from commercial suppliers and were used as received unless otherwise indicated. Analytical thin-layer chromatography (TLC) on silica gel 60F254 aluminum precoated plates (0.25 mm layer) was used for monitoring reaction progress. The components were visualized using UV light at 254 nm. All chromatographic purifications were performed using flash column chromatography on silica gel (200–400 mesh). The solvents were removed by rotary evaporation using a Rotathe authors’Detailsvapor. ^1^H and ^13^C NMR (δ in ppm) spectra (Supplementary materials Data S3-S14) were recorded in CDCl_3_ (unless otherwise noted) at 500 and 125.8 MHz, respectively (unless otherwise noted). Residual CHCl_3_ (δ = 7.26) as a reference for 1H NMR spectra and carbon absorption of CDCl3 (δ = 77.0) as an internal reference for ^13^C NMR spectra were used. Data are reported as s, singlet; d, doublet; t, triplet; q, quartet; dd, doublet of doublets; dt, doublet of triplets; td, triplet of doublets; tt, triplet of triplets; m, multiplet. Infrared (IR) spectra were recorded using an ATR instrument in neat samples. High-resolution mass spectra (HR-MS) were acquired at the Shimadzu Center for Advanced Analytical Chemistry using electrospray ionization (ESI), with mass measured using electrospray ionization (ESI-TOF).

#### Synthesis of Thiourea derivatives

General procedure for the preparation of thioureas derived from 1-(4-methoxyphenyl)-*N*-methylmethanamine: To a solution of 1-(4-methoxyphenyl)-*N*-methylmethanamine (3.31 mmol, 0.51 ml, 1.0 equiv) in DCM (30 ml) was added triethylamine (3.97 mmol, 0.55 ml, 1.2 equiv) followed by stirring the reaction mixture for 15 minutes. The corresponding aryl isothiocyanate was added (3.31 mmol, 1.0 equiv) at rt and stirring was continued overnight at rt. The solvent was evaporated, and the crude thioureas were purified by chromatography.

#### 1-(4-Methoxybenzyl)-1-methyl-3-phenylthiourea (6a)

Obtained as a white solid (0.93 g, 98%), m.p. 126-129 °C, purified by chromatography (25% EtOAc/Hexanes). ^1^H NMR (500 MHz, CDCl_3_) δ 7.37 – 7.08 (m, 7H), 6.96 – 6.82 (m, 2H), 5.01 (s, 2H), 3.81 (s, 3H), 3.25 (s, 3H); ^13^C NMR (125 MHz, CDCl_3_) δ 182.6, 159.4, 139.8, 128.8, 128.7, 127.9, 125.8, 125.3, 114.4, 56.6, 55.4, 38.9; FT-IR (neat, cm^-1^): 3260, 3042, 2996, 2924, 2831, 1610, 1592; HRMS (*m/z*): calc for [M+H]^+^ C_16_H_178_N_2_O_S_ 287.1152 found 287.1190.

#### 1-(4-Methoxybenzyl)-3-(4-methoxyphenyl)-1-methylthiourea (6b)

Obtained as white solid (0.95 g, 90%), m.p. 108-110 °C, purified chromatography (15% EtOAc/Hexanes). ^1^H NMR (500 MHz, CDCl_3_) δ 7.25 – 7.18 (m, 2H), 7.15 – 7.05 (m, 3H), 6.89 – 6.76 (m, 4H), 4.96 (s, 2H), 3.75 (s, 3H), 3.73 (s, 3H), 3.19 (s, 3H). ^13^C NMR (125 MHz, CDCl_3_) δ 182.9, 159.4, 157.9, 132.7, 128.7, 128.0, 127.8, 114.4, 114.0, 56.5, 55.5, 55.4. FT-IR (neat, cm^-1^): 3315, 3010, 2950, 2916, 2831, 1610, 1594; HR-MS (*m/z*): calc for [M+H]^+^ C_17_H_20_N_2_O_2_S 317.1318 found 317.1302.

#### 1-(4-Methoxybenzyl)-1-methyl-3-(p-tolyl)thiourea (6c)

Obtained as a white solid (0.89 g, 90%), m.p. 142-144 °C, purified chromatography (15% EtOAc/Hexanes). ^1^H NMR (500 MHz, CDCl_3_) δ 7.28 – 7.22 (m, 3H), 7.18 – 7.09 (m, 4H), 6.93 – 6.86 (m, 2H), 4.99 (s, 2H), 3.80 (s, 3H), 3.15 (s, 3H), 2.33 (s, 3H). ^13^C NMR (126 MHz, CDCl_3_) δ 182.6, 159.3, 137.3, 135.7, 129.3, 128.8, 128.2, 126.0, 114.3, 56.5, 55.4, 29.8, 21.1. FT-IR (neat, cm^-1^): 3324, 3261, 2999, 2918, 2834, 1611, 1589, 1510; HR-MS (*m/z*): calc for [M+H]^+^ C_17_H_20_N_2_OS 301.1369 found 301.1353.

#### 3-(4-Chlorophenyl)-1-(4-methoxybenzyl)-1-methylthiourea (6d)

Obtained as yellow solid (0.9 g, 85%), m.p. 148-150 °C, purified chromatography (30% EtOAc/Hexanes). ^1^H NMR (500 MHz, CDCl_3_) δ 7.36 – 7.11 (m, 7H), 6.98 – 6.84 (m, 2H), 5.00 (s, 2H), 3.82 (s, 3H), 3.23 (s, 3H). ^13^C NMR (125 MHz, CDCl_3_) δ 182.3, 159.4, 138.4, 131.1, 128.7, 128.7, 127.8, 127.0, 114.4, 56.5, 55.4. FT-IR (neat, cm^-1^): 3235, 3036, 3010, 2963, 2926, 2781, 1610, 1587, 1510, 1325; HR-MS (*m/z*): calc for [M+H]^+^ C_16_H_17_ClN_2_OS 321.0823 found 321.0788.

#### 3-(3-Chloro-4-methylphenyl)-1-(4-methoxybenzyl)-1-methylthiourea (6e)

obtained as a white solid (0.78 g, 71%), m.p. 146-148 °C, purified by ethanolic recrystallization and dried at room temperature. ^1^H NMR (500 MHz, CDCl_3_) δ 7.24 – 7.13 (m, 4H), 7.08 (d, *J* = 8.0 Hz, 1H), 7.01 (dt, *J* = 8.2, 1.8 Hz, 1H), 6.86 – 6.80 (m, 2H), 4.92 (s, 2H), 3.74 (s, 3H), 3.12 (s, 3H), 2.26 (s, 3H). ^13^C NMR (126 MHz, CDCl3) δ 182.4, 159.4, 138.6, 133.9, 133.6, 130.7, 128.7, 127.9, 126.2, 124.4, 114.4, 56.6, 55.4, 38.6, 19.7. FT-IR (neat, cm^-1^): 3230, 3083, 3000, 2956, 2829, 1577, 1509, 1491, 1395, 882; HR-MS (*m/z*): calc for [M+H]^+^ C_16_H_20_ClN_2_OS 335.0979 found 335.0957

#### Synthesis of benzothiazole derivatives

The thiourea (1.0 equiv) was dissolved in HFIP (25 mL), then Cs_2_CO_3_ (1.2 equiv) was added and stirred for 10 min. 1.0 or 2.0 equiv of IBDA was added, followed by stirring the reaction mixture for 2 h. The reaction was quenched with saturated brine solution (10 ml), then extracted with dichloromethane (3 × 10 ml), and dried over sodium sulfate. The crude benzothiazoles were obtained by removing the organic solvent by rotary evaporation and subjected to chromatographic purification.

#### *N*-(4-Methoxybenzyl)-*N*-methylbenzo[*d*]thiazol-2-amine (7a)

Obtained via oxidative dearomatization of 1-(4-methoxybenzyl)-1-methyl-3-phenylthiourea (**6a**) as a white solid (9%, 18.0 mg, using 2.0 equiv PIDA), (47%, 220 mg, using 1.0 equiv IBDA), m.p. 74-76 °C, purified by chromatography (1% EtOAc/Hexanes). ^1^H NMR (500 MHz, CDCl_3_) δ 7.58 (ddd, *J* = 7.7, 7.2, 1.2 Hz, 2H), 7.30 (ddd, *J* = 7.9, 7.3, 1.3 Hz, 1H), 7.27 – 7.20 (m, 2H), 7.06 (td, *J* = 7.5, 1.2 Hz, 1H), 6.89 – 6.84 (m, 2H), 4.70 (s, 2H), 3.79 (s, 3H), 3.11 (s, 3H); ^13^C NMR (125 MHz, CDCl_3_) δ 168.9, 159.3, 153.3, 131.1, 129.1, 128.5, 126.1, 121.2, 120.7, 118.9, 114.27, 114.2, 56.1, 55.4, 37. 7, 29; FT-IR (neat, cm^-1^): 196 2960, 2917, 2853, 2832, 1596, 1543, 1410; HR-MS (*m/z*): calc for [M+H]^+^ C_16_H_16_N_2_OS 285.1056, found 285.1045.

#### 2-((4-Methoxybenzyl)(methyl)amino)benzo[d]thiazol-6-yl acetate (7b)

Obtained via oxidative dearomatization of 1-(4-methoxybenzyl)-1-methyl-3-phenylthiourea (**6a**) as a white solid (27%, 66.0 mg, using 2.0 equiv IBDA), m.p. 100-102 °C, purified by chromatography (8% EtOAc/Hexanes). ^1^H NMR (500 MHz, CDCl_3_) δ 7.52 (d, J = 8.7 Hz, 1H), 7.34 (d, J = 2.4 Hz, 1H), 7.22 (d, *J* = 8.7 Hz, 2H), 7.00 (dd, *J* = 8.7, 2.4 Hz, 1H), 6.88 – 6.84 (m, 2H), 4.68 (s, 2H), 3.79 (s, 3H), 3.09 (s, 3H), 2.30 (s, 3H); ^13^C NMR (126 MHz, CDCl_3_) δ 170.1, 169.0, 159.4, 151.2, 144.8, 131.5, 129.1, 128.3, 119.8, 119.1, 114.2, 113.9, 56.1, 55.4, 37.7, 29.8, 21.2; FT-IR (neat, cm^-1^): 2920, 2850, 1758, 1602, 1574, 1509; HR-MS (*m/z*): calc for [M+H]^+^ C_16_H_16_N_2_OS 285.1056, found 285.1045.

#### 6-Methoxy-N-(4-methoxybenzyl)-N-methylbenzo[d]thiazol-2-amine (7c)

Obtained as white solid (23%, 70.0 mg, using 2.0 equiv PIDA), (65%, 0.321 g, using 1.0 equiv IBDA), m.p. 86-88 °C, purified chromatography (10% EtOAc/Hexanes). ^1^H NMR (500 MHz, CDCl3) δ 7.59 (d, *J* = 8.8 Hz, 1H), 7.41 – 7.32 (m, 2H), 7.26 (d, *J* = 2.6 Hz, 1H), 7.08 – 6.94 (m, 3H), 4.78 (s, 2H), 3.93 (s, 3H), 3.91 (s, 3H), 3.20 (s, 3H). ^13^C NMR (125 MHz, CDCl_3_) δ 167.5, 159.3, 154.9, 147.5, 132.0, 129.1, 128.6, 119.3, 114.2, 113.6, 105.4, 56.1, 56.0, 55.4, 37.6, 29.8. FT-IR (neat, cm^-1^): 2920, 2854, 1605, 1546, 1508, 1211; HR-MS (*m/z*): calc for [M+H]^+^ C_17_H_18_N_2_O_2_S 315.1162 found 315.1153.

#### *N*-(4-Methoxybenzyl)-*N*,6-dimethylbenzo[*d*]thiazol-2-amine (7e)

Obtained as a white solid (20%, using 2.0 equiv IBDA), m.p. 80-82 °C, purified chromatography (25% EtOAc/Hexanes). ^1^H NMR (500 MHz, CDCl_3_) δ 7.44 (d, *J* = 8.2 Hz, 1H), 7.36 – 7.32 (m, 1H), 7.22 – 7.15 (m, 2H), 7.11 – 7.04 (m, 1H), 6.85 – 6.77 (m, 2H), 4.63 (s, 2H), 3.74 (s, 3H), 3.05 (s, 3H), 2.35 (s, 3H). ^13^C NMR (125 MHz, CDCl_3_) δ 182.6, 159.3, 137.3, 135.7, 129.3, 128.8, 128.2, 126.0, 114.3, 56.5, 55.4, 38.2, 21.1. FT-IR (neat, cm^-1^): 2994, 2924, 2850, 1629, 1602, 1494; HR-MS (*m/z*): calc for [M+H]^+^ C_17_H_18_N_2_OS 299.1213 found 299.1194.

#### 6-Chloro-N-(4-methoxybenzyl)-N-methylbenzo[d]thiazol-2-amine (7f)

Obtained as a white solid (60 mg, 30%, using 1.0 equiv IBDA), m.p. 72-73 °C, purified chromatography (8% EtOAc/Hexanes). ^1^H NMR (500 MHz, CDCl_3_) δ 7.54 (dd, *J* = 2.2, 1.0 Hz, 1H), 7.47 (dd, *J* = 8.6, 1.0 Hz, 1H), 7.28 – 7.20 (m, 3H), 6.92 – 6.83 (m, 2H), 4.68 (s, 2H), 3.79 (s, 3H), 3.10 (s, 3H). ^13^C NMR (125 MHz, CDCl_3_) δ 168.9, 159.4, 151.6, 132.1, 129.1, 128.1, 126.5, 126.2, 120.4, 119.5, 114.2, 56.1, 55.3, 37.7. FT-IR (neat, cm^-1^): 3010, 2961, 2924, 2836, 1593, 1560, 1509, 1448, 1412; HR-MS (*m/z*): calc for [M+H]^+^ C_16_H_16_ClN_2_OS 319.0650 found 319.0666.

#### 5-Chloro-N-(4-methoxybenzyl)-N,6-dimethylbenzo[d]thiazol-2-amine (7g)

Obtained as a white solid (44 mg, 22%, using 1.0 equiv IBDA), m.p. 104-106 °C, purified chromatography (5% EtOAc/Hexanes). ^1^H NMR (500 MHz, CDCl_3_) δ 7.52 (s, 1H), 7.35 (s, 1H), 7.28 – 7.10 (m, 2H), 6.87 – 6.76 (m, 2H), 4.63 (s, 2H), 3.75 (s, 3H), 3.04 (s, 3H), 2.34 (s, 3H). ^13^C NMR (126 MHz, CDCl_3_) δ 169.2, 159.3, 132.3, 130.2, 129.1, 128.6, 128.1, 121.9, 119.0, 114.2, 114.0, 56.1, 55.3, 37.7, 20.1. FT-IR (neat, cm^-1^): 3056, 3032, 2990, 2918, 2849, 2834, 1737, 1606, 1561, 1176, 1098, 868; HR-MS (*m/z*): calc for [M+H]^+^ C_17_H_18_ClN_2_OS 333.0823 found 333.0799.

### 2.3 C2C12 Cell Culture and In-Vitro Studies

#### 2.3.1 Cell’s Viability using MTS Cytotoxicity assay

All cell culture studies were performed according to our published protocols. (*64*) C2C12 skeletal myoblast cells were seeded in 24-well plates with 50% confluency and incubated for 6 hours to allow the cells to settle down and attach. After 6 hours, the growth media was removed, and 500 µL of conditioned media was added to each well. Conditioned media contains DMEM, 10% FBS, and 2% P/S with a specific concentration of each compound. Five different concentrations of each compound were used: 5 µM, 10 µM, 50 µM, 75 µM, and 100 µM. The used benzothiazole compounds were dissolved in 1mL of DMSO, followed by serial dilution in cell culture media to have the exact final concentrations of each tested compound. DMSO percentage did not exceed 0.1% in the final media used, and the precise concentrations of DMSO were used as a control. Cells seeded in a tissue culture plate with a normal growth medium were used as a positive control for each group. Exact concentrations of DMSO were tested in a separate plate to reveal any effect that might be introduced due to using DMSO as a vehicle for the tested compounds. After 24 hours, the conditioned media were collected and stored at -20 °C for possible further analysis. The MTS assay kit was used to quantify the cells’ viability/cytotoxicity of each tested compound, and each group was tested with n = 4. After adding 100 µL of the MTS reagent, plates were wrapped in aluminum foil to avoid direct light and incubated for 2 hours in a 5% CO_2_ and 37 °C incubator. Then, the optical density (OD) readings were recorded using a microplate reader (SpectraMax^®^ i3, Molecular Devices, CA, USA) at 490 nm. Data are presented as box plots with whiskers presenting the standard deviation (SD), and the (±) indicates the mean of each group.

#### 2.3.2 Cell Differentiation and Myogenic Studies

C2C12 cells were cultured for 5 days in differentiation media (DM) to study the effect of the benzothiazole agents on cell morphology and differentiation capacity. For each benzothiazole agent, 2 different concentrations (5 μM and 10 μM) were dissolved in DMSO and used for the differentiation study compared to normal DM as controls. The final DMSO concentration in each well was 0.05% and 0.1% for 5 µM and 10 μM, respectively. Thus, the same concentrations of DMSO were tested to capture any effect that might be introduced due to the use of DMSO as a solvent/vehicle for the tested compounds. DMSO 0.05% and 0.1% were labeled as DMSO-5 μM and DMSO-10 μM, respectively, to compare the agent concentrations easily. 5 × 10^4^ cells/well were seeded in 12-well plates in GM for all differentiation experiments until 75% confluency. GM was removed after reaching 75% confluency, and each well was washed with 2 mL of PBS-1× to remove any unattached cells. Then, 1 mL of condition media (DM+5 μM and 10 μM of each compound) was added to each well and compared to normal DM and agents-free DMSO (0.05% and 0.1%) as controls. DM was collected for biomarker expression assays, and new DM with the specified agent concentration was added to each well every 48 hours. After 5 days, cells were fixed and immunostained following previously published protocols.(*64*) Briefly, DM was removed, and cells were washed 3 times with PBS-1×, then fixed with 2 mL of neutral buffered formalin for 7–8 min. Cells were rewashed and permeabilized with 2 mL of 0.1% triton X-100 in PBS for 10 min. Then, cells were stained with 20 µL/mL of conjugated MHC antibody in 1× TBST (PBS with 0.1% Triton and 0.1% Tween) for 45 min and counterstained with DAPI (1 ug/µL). Finally, cells were washed with 2 mL/well of PBS-1× for 5 min, 3 times, and fluorescence images were captured at 10× or 20× using the Leica DMi8 Inverted Fluorescence Microscope.

#### 2.3.3 Quantification of aminobutyric acids

To investigate the altered levels of GABA and its isomers generated by C2C12 cells after the treatment of different concentrations of benzothiazole agents **7b** and **7g**, an LC-MS/MS-based analytical method was performed to quantify GABA, α-aminobutyric acid (AABA, L- and D-), β-aminobutyric acid (BABA, L), and β-aminoisobutyric acid (BAIBA, L- and D-) in cells and conditioned media (CM) (PMID 31969651). The LC-MS/MS analysis was performed on a Shimadzu LCMS-8050 triple quadrupole mass spectrometer (Shimadzu Scientific Instruments Inc., Tokyo, Japan). LC separation was conducted on a chiral SPP-TeicoShell column (150 × 4.6 mm, 2.7 µm, AZYP LLC., Arlington, TX) configured with a Max-RP column (50 × 2.0 mm, Phenomenex, Torrance, CA). The mobile phase consists of (A) methanol and (B) 0.005% formic acid and 2.5 mM ammonium formate in water. The instrument was operated and optimized using pure standard solutions under positive electrospray ionization (ESI) and multiple reaction monitoring (MRM) modes. Ten microliter CM samples and same volume of isotopic internal standards (IS) mixture solution (1.2 µM of D, L-BAIBA-d_3_, D, L-AABA-d_6_, and GABA-d_6_, 0.1% formic acid in methanol, v/v) were added to 35 µL of 0.1% formic acid in methanol (v/v), followed by 20 min-shaking at room temperature and then 15 min-centrifugation at 15,000 ×g, 4 °C to precipitate protein. The supernatant was directly transferred to an autosampler vial, and 45 μL of each sample was injected for LC-MS/MS analysis. All analyses and data processing were completed on Shimadzu LabSolutions V5.91 software.

#### 2.3.4 RNA isolation and RT-PCR gene arrays

Total RNA from C2C12 cells on the 5^th^ day post-treatment with 10 µM benzothiazole agents **7b** and **7g** and vehicles (DMSO) were extracted from the cells with TRI reagent according to the manufacturer’s protocol. We used the Mouse GABA & Glutamate RT² Profiler PCR Array to detect gene expression changes in cells treated with benzothiazole agents **7b** and **7g**. cDNA was synthesized using the RT^2^ First Strand Kit (this kit contains genomic DNA elimination buffer to eliminate genomic DNA), and the PCR Array was run according to the manufacturer’s instructions, including a threshold of 0.2. Data were analyzed using the RT^2^ Profiler™ PCR Array Data Analysis Software; Ct values were normalized to six built-in reference housekeeping genes, genomic DNA control, reverse transcription control, and positive PCR control. We used this analytical software (https://geneglobe.qiagen.com/us/analyze/) to set the significance of up/downregulation of all tested genes at a two-fold difference and p < 0.05. Three independent experiments were performed with a sample size of n = 3 per group.

#### 2.3.5 Reactive Oxygen Species Detection and Quantification

C2C12 cells were grown in 24-well plates until 70% confluency in a standard growth medium. Then, 8 wells were incubated with 10 µM of benzothiazole agents **7b** and **7g** for 30 minutes as a pre-treated group. After, cells were washed with PBS, and all wells were incubated with 10 µM of carboxy-H2DCFDA (D399, Invitrogen, CA, USA) for 15 minutes at 37 °C in the dark. Then, cells were washed 3 times with PBS and treated with growth media containing different treatments. Regular media as a positive control, normal media+100 µM H_2_O_2_ as a negative control, regular media for the pre-treated group, and everyday media+10 µM of agents **7b** and **7g** as co-treated groups. Again, cells were incubated for 30 minutes at 37 °C in the dark and then imaged using a fluorescent microscope. ImageJ software was used to analyze the fluorescent images, and the results are presented as ROS fold change in relative fluorescence.

#### 2.3.6 Lipidomics analysis

C2C12 cells were incubated with benzothiazoles **7b** and **7g** for 24 hours, then culture media was removed, and cells were washed with fresh media and then centrifuged to isolate the cell pallets. The cells were homogenized and prepared following the published solid-phase extraction protocol(*65*). The final samples were recovered in methanol and then concentrated using a centrifugal vacuum concentrator. The lipidomics analysis was performed using an optimized LC/ESI-MS/MS method utilizing the Shimadzu LCMS-8060 system (Shimadzu Scientific Instruments, Inc., Columbia, MD, USA) to profile and quantify a total of 158 lipid mediators. The mobile phase used was 0.1% Formic acid in water (Pump A), and 100% Acetonitrile (Pump B). The LC system is equipped with a system controller: CBM-20A, three LC-30AD pumps, a SIL-30AC autosampler, and a CTO-30A column oven. The LC separation is conducted on a C8 column (Ultra C8, 150×2.1mm, 3μm, RESTEK, Manchaca, TX, USA). The oven temperature for the analytical column is set to 40°C. The triple quadrupole mass spectrometer (LCMS-8060 system) is operated and optimized under electrospray ionization in negative mode. MS parameters are optimized as follows: interface: DUIS (ESI) and (APCI), voltage, 4.0kV; interface temperature, 280°C; desolvation line (DL) temperature, 275°C; heating block temperature, 400°C; drying gas (N2), 10L/min; nebulizing gas (N2), 3L/min; heating gas (air), 10L/min; and collision-induced dissociation (CID) gas (Ar), 230 kPa. Once all parameters have been optimized, the optimized method is run with a full range to scan and monitor all internal standards (IS) and standard lipid mediators (Std) in a concentrated sample of IS + Std mixture. Once all parameters are optimized and the calibration curve results meet the standards, samples will be loaded. The samples prepared by solid phase extraction are reconstituted with the dried extracts in the 1.5 mL tubes by adding 50μL of methanol, vortexing, and then transferring 20μL of each sample directly into autosampler glass vials for auto injection of 10μL. After all samples are run successfully, the raw data are uploaded into the LabSolutions software, along with the calibration curve results, to quantify the detected LMs in the tested samples.

#### 2.3.7 Qiagen Ingenuity Pathway Analysis (IPA)

IPA analyses were performed using the North Texas Genomics Center (NTGC) multi-user license (Qiagen Ingenuity IPA version 145030503) for the IPA software, with direct support from a QIAGEN Field Application Scientist. The IPA software offers capabilities for pathway analysis, causal network modeling, data integration, and interactive visualization. The IPA is a comprehensive bioinformatics solution for the analysis and interpretation of ‘omics data within a biological and disease context. Its capabilities are widely used in genomics, transcriptomics, proteomics, and metabolomics research to extract meaningful biological insights from complex datasets. Using IPA, we predict upstream regulators, downstream effects, and help generate causal hypotheses about gene expression changes. The platform integrates a vast, expertly curated knowledge base that connects genes, proteins, chemicals, and diseases for deep biological insight.

### 2.4 Statistical Analysis

All experiments were conducted with three or more independent biological replicates (n ≥ 3) unless specified otherwise. Data are presented as mean ± standard deviation (SD) or box plots with whiskers representing SD; individual means are indicated by ± symbols. Statistical analyses were performed using OriginPro software (Version 2025, OriginLab Corporation, Northampton, MA, USA). For comparison among multiple groups, one-way analysis of variance (ANOVA) followed by Tukey’s post hoc test was used. For comparisons between two groups, unpaired two-tailed Student’s T-tests were applied. Gene expression data from PCR arrays were analyzed using Qiagen’s RT² Profiler PCR Array Data Analysis software; significance was defined as fold change ≥ 2.0 with p < 0.05. Statistical significance for all experiments was set at p < 0.05, and exact statistical tests used for each dataset are specified in figure legends.

## 3. RESULT

### 3.1 Organic Synthesis

Most reported methods for synthesizing benzothiazole derivatives require relatively harsh reaction conditions, at elevated temperatures, metal-catalyzed transformations, and long completion times. In this study, we report a rapid and straightforward approach to the benzothiazole scaffold through environmentally benign conditions developed by our group for this chemoselective cyclization using a hypervalent iodine reagent. While investigating the dearomatization reactions of benzylic thioureas, we discovered that some substrates afforded benzothiazole rather than the expected cyclohexadienone. Selective oxidation of the precursor thiourea containing two aryl groups **6a-e** (**Scheme 1**) derived from the corresponding isothiocyanate occurs presumably through activation of sulfur, followed by electrophilic aromatic substitution of the N-aryl group rather than addition to the N-benzyl group. However, the present methodology requires further optimization to identify its limitations. It offers a straightforward approach for constructing benzothiazole derivatives from simple thioureas, thereby delivering the substituted core structure of the benzothiazole scaffold. Subjecting the synthesized thioureas (**6a-e**) to iodosobenzene diacetate (IBDA) in the presence of Cs_2_CO_3_ in hexafluoroisopropanol (HFIP) as a solvent yielded the corresponding benzothiazoles, as shown in **Scheme 1**. Interestingly, the chemoselectivity of the formation of acetylated benzothiazole depends mainly on the molar equivalents of IBDA (**Table 1**). Therefore, two acetylated benzothiazoles (**7b** and **7d**) were isolated when a 2.0 equivalent of IBDA was used. The X-ray crystal structure of **7b** confirmed the chemical structures (CCDC: 2034523)(*66*) in addition to other spectroscopic techniques. The initial optimization study was performed by lowering the IBDA equivalents to 1 molar equivalent, yielding good-to-moderate yields of benzothiazoles **7a** and **7c**.

**Scheme 1.**
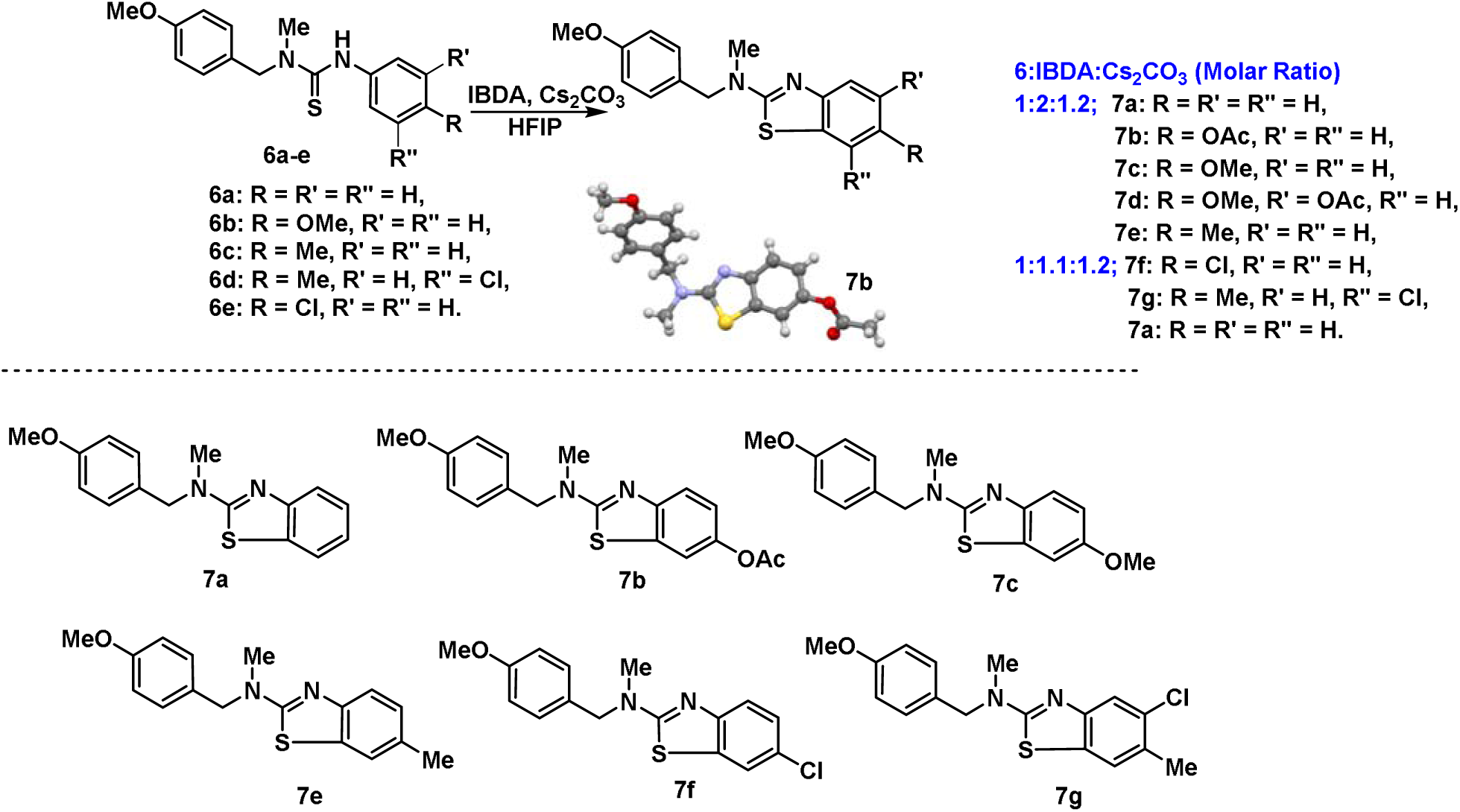
Oxidation electrophilic aromatic substitution of thioureas **6a**-**e** towards targeted 2-aminobenzothiazole derivatives, resulting in the isolation of benzothiazoles **7a-g**.

**Table 1.** Reaction conditions and yields of the isolated benzothiazoles (6a-7g)

|  | Thiourea derivatives | PIDA equivalents | Benzothiazoles |  |  |  | Yield (%) |
| --- | --- | --- | --- | --- | --- | --- | --- |
|  |  |  | Code | R | R' | R'' |  |
| **1** | 6a | 1.0 | 7a | H | H | H | 47 |
| **2** | 6a | 2.0 | 7a | H | H | H | 11 |
| **3** | 6a | 2.0 | 7b | OAc | H | H | 27 |
| **4** | 6b | 1.0 | 7c | OMe | H | H | 65 |
| **5** | 6b | 2.0 | 7c | OMe | H | H | 23 |
| **6** | 6b | 2.0 | 7d | OMe | H | OAc | 14 |
| **7** | 6c | 1.0 | 7e | Me | H | H | 19 |
| **8** | 6c | 2.0 | 7e | Me | H | H | 20 |
| **9** | 6d | 1.0 | 7f | Cl | H | H | 30 |
| **10** | 6e | 1.0 | 7g | Me | Cl | H | 22 |

### 3.2 Biological studies of benzothiazoles

#### Cytotoxicity of benzothiazoles

To study the cytotoxicity of benzothiazoles, C2C12 skeletal muscle cells were cultured, and the cells’ viability was assessed using an MTS cell viability assay. C2C12 cell viability was measured after 24 hours of proliferation and treatment with various concentrations of each tested benzothiazole agent (5, 10, 50, 75, and 100 µM). All results are normalized to dimethyl sulfoxide (DMSO), used as the vehicle, and compared with a positive control (normal growth medium), as shown in **Figure 2**. All tested compounds showed no cytotoxic effect at lower doses (5 and 10 µM), whereas significant decreases in C2C12 cell viability were observed at higher concentrations (50, 75, and 100 µM) for all compounds. Furthermore, low concentrations of agents **7b**, **7c**, **7e**, and **7g** significantly increased C2C12 cell viability after 24 hours of proliferation, indicating a beneficial effect on cell proliferation.

**Figure 2:**
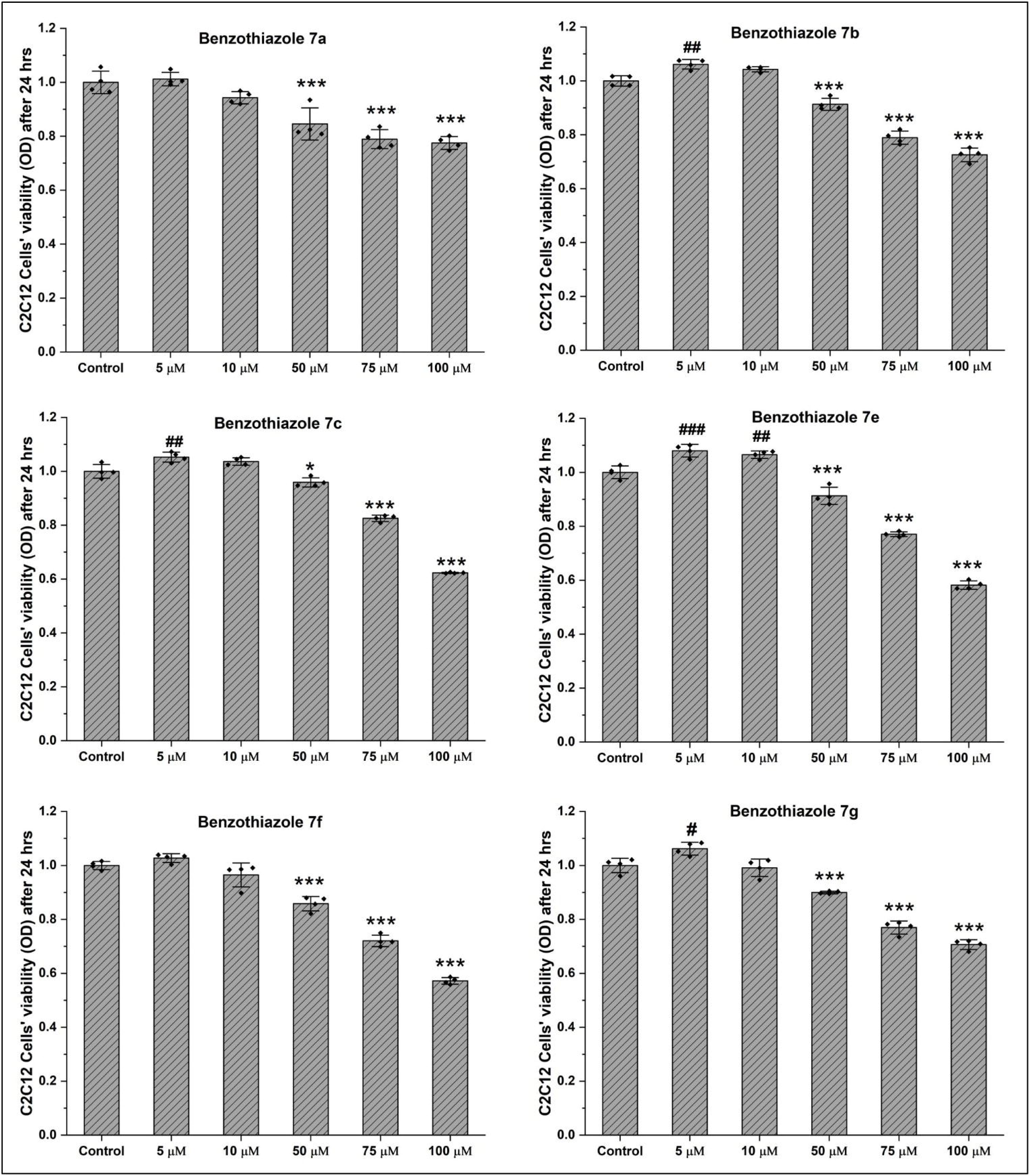
Cytotoxicity of benzothiazoles using MTS assay. Six different compounds of benzothiazole derivatives were screened at different concentrations to test their cytotoxicity effect on C2C12 cells. Low concentrations of 5 µM to 10 µM showed no cytotoxicity while higher doses indicated a significant decrease in cell viability, as shown *p<0.05, **p<0.01, ***p<0.001. Compounds **7b**, **7c**, **7e**, and **7g** indicated a significant increase in cell viability at a low dose of 5µM, as shown # p<0.05, ## p<0.01, ###p<0.001.

#### Myogenic effects of benzothiazoles

To study the myogenic effects of the benzothiazoles, C2C12 skeletal muscle cells were cultured and allowed to differentiate for 5 days in differentiation medium supplemented with 5 μM and 10 μM solutions of each benzothiazole agent in DMSO, as shown in **Figure 3**. Figure 3-A shows the differentiated C2C12 cells (myotubes) stained with Myosin Heavy Chain (MHC-green) antibodies and 4′,6-diamidino-2-phenylindole dihydrochloride hydrate (DAPI-blue), which stains the nuclei. Figure 3-B depicts the quantitative analysis of the differentiation study. Bar graphs showing the distribution of data points show the mean values of all agents’ fusion index (FI) compared to the control and the vehicle (DMSO, 5-10 µM). FI is considered a standard method for measuring myotubule formation in muscle cells and quantifying the differentiation phase. It measures the ratio of fused nuclei and formed myotubes divided by the total number of nuclei. Thus, FI indicates the ability of mononucleated single cells to fuse and form multinucleated myotubes. Analysis showed that the 5 µM DMSO groups had no significant effect on FI compared with the positive control, whereas DMSO (10 µM) slightly increased the FI. A low dose of benzothiazole agents did not show any significant changes in the FI compared to the control, except for **7b** (5 µM), which induced a significant decrease in the FI compared to the positive control. The high concentration of 10 µM for all benzothiazole agents resulted in a substantial reduction in FI compared to the control, except for agent **7g**, which showed an FI ratio similar to the positive control at both 5 and 10 µM. It is important to note that only agent **7b** presented a significant decrease in FI at both concentrations. In contrast, agent **7g** did not show a significant difference, even at high concentrations. Thus, benzothiazoles **7b** and **7g** were selected for further omics and RT-PCR analysis.

**Figure 3:**
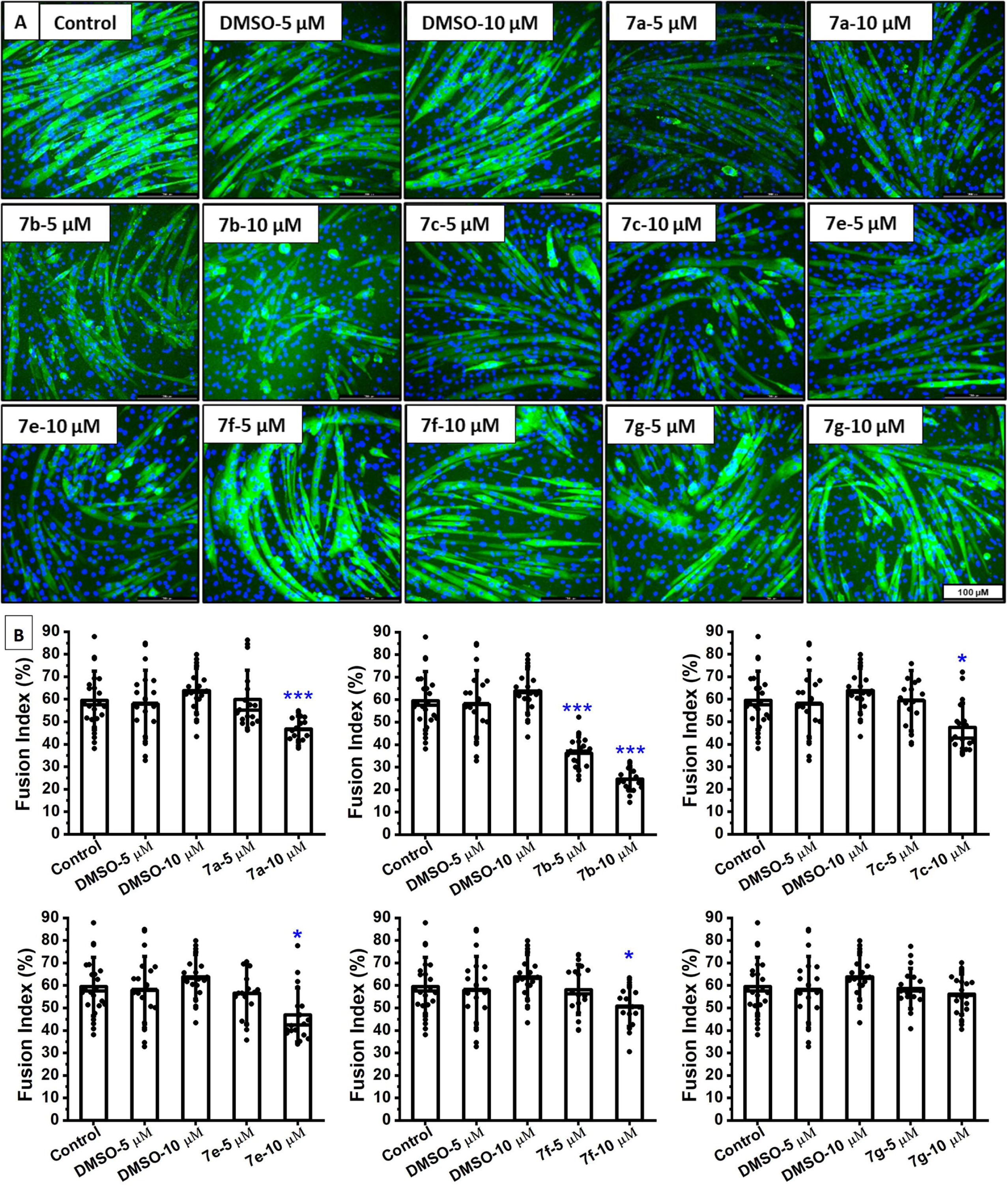
Myogenic effects of benzothiazoles. Six different compounds of benzothiazoles derivatives were studied at two concentrations to test their myogenic effect on C2C12 cells. A) Differentiated C2C12 cells (Myotubes) stained with Myosin Heavy Chain and DAPI. B) Calculated fusion index of tested benzothiazoles compared to positive control and DMSO, statistical analysis based on One-Way ANOVA with Tukey Post Hoc test; * p<0.05, **p<0.01, ***p<0.001.

#### Effect of benzothiazoles on GABA

To investigate the effect of benzothiazoles on GABA and its isomers generated in C2C12 cells and in conditioned media after treatment with different concentrations of benzothiazole agents, our novel LC-MS/MS-based method was used (*67, 68*). GABA, α-aminobutyric acid (*L- and D-*AABA), β-aminobutyric acid (*L-*BABA), and β-aminoisobutyric acid (*L- and D-*BAIBA) were measured and quantified in cells and conditioned media (CM). GABA was determined in C2C12 cells treated with the vehicle (DMSO), control cells, and cells treated with 10 µM benzothiazole agents **7b** and **7g**. At the same time, L-BAIBA was detected only in cells treated with benzothiazoles, as shown in **Figures 4A and 4B**. Treating cells with benzothiazoles reduced GABA concentrations compared with the positive control and the DMSO vehicle. Benzothiazole **7b** significantly decreased the GABA concentration (p < 0.05), while agent **7g** showed no significant difference. On the other hand, only cells treated with agent **7b** showed a substantial increase in the L-BAIBA concentration compared to the control and the DMSO vehicle (p < 0.05). Furthermore, GABA, L-BAIBA, and D-BAIBA were determined in the media of C2C12 cells on the 5^th^ day post-treatment with 10 µM benzothiazole agents **7b**, **7g**, and vehicle (**Figure 4C-E**). GABA was generated by C2C12 and released into the surrounding media (**Figure 4C**). After treatment with **7b** or **7g** (10 µM), GABA concentration in CM was significantly lower than that found in the control and DMSO-5 µM groups, but this reduction was not significant when compared with the DMSO-10 µM group (**Figure 4C**). A significantly higher L-BAIBA level was observed in CM on the 5^th^ day when C2C12 were treated with 10 µM of **7b** compared to all other groups, including control, DMSO vehicle, and **7g** (**Figure 4D**). No significant alteration was observed in the D-BAIBA concentrations in CM (**Figure 4E**). These omics data indicated that the benzothiazole agent **7b** significantly decreased GABA levels and increased L-BAIBA concentrations and secretion in C2C12 cells and in the media. Additionally, benzothiazole **7g** slightly decreased the GABA concentration in cells and did not significantly alter the concentrations of L- and D-BAIBA. Statistical analysis using One-way ANOVA with Tukey’s post-hoc test (α = 0.05) was performed for multiple comparisons between groups in this study.

**Figure 4:**
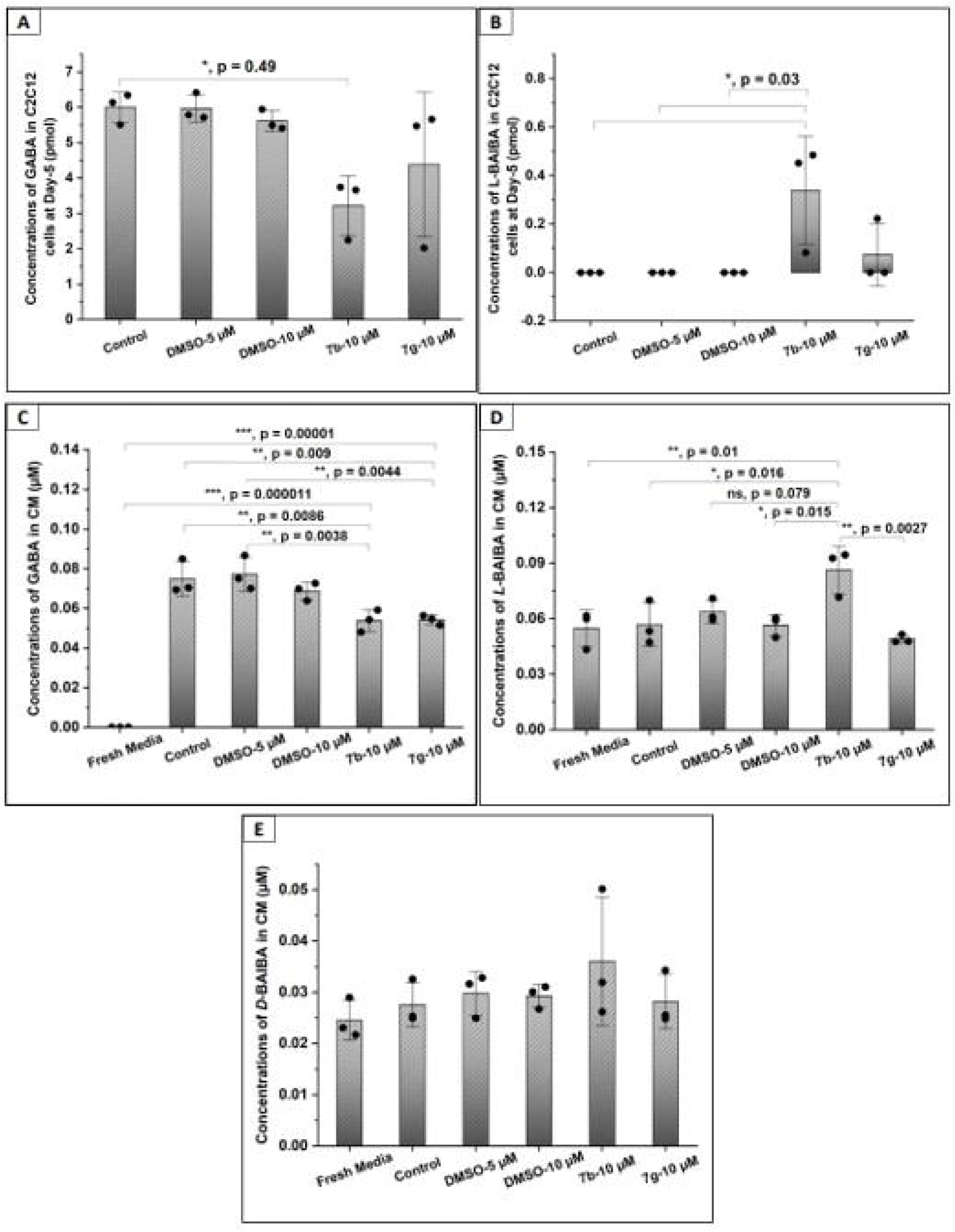
Concentrations of aminobutyric acids in conditioned media (CM) and C2C12 cells at the 5^th^ day post-treatment with 10 µM benzothiazole agents 7b and 7g. (A) GABA in cells, (B) *L*-BAIBA in cells, (C) GABA in CM, (D) L-BAIBA in CM, and (E) *D*-BAIBA in CM. Mean ± SD (n=3). One-way ANOVA with Tukey’s post-hoc test (α = 0.05) was performed for multiple comparisons between groups.

#### PCR Gene Array

Based on the omics results, a GABA RT² Profiler PCR Gene Array was performed to gain further insights into the effect of benzothiazoles on GABA gene regulation. For the GABA gene array experiments, differentiated C2C12 cells were treated with 10 µM of benzothiazole agents **7b** and **7g** compared to control and 10 µM DMSO. Then, the mouse GABA and Glutamate RT² Profiler PCR Array was performed. This mouse GABA and glutamate profiler PCR array profiles 84 key genes of the GABA and glutamate neurotransmitter system. The results of this array indicated that benzothiazole **7b** significantly decreased the expression of GABA receptor Gabrg2 (Fold change is -2.25, p = 0.016) compared to DMSO 10 µM. Furthermore, agent **7b** reduced the expression of the neurotransmitter receptors Gria1, Grin2a, and Grm7 (Fold changes are -2.26, -2.33, and -2.79, respectively) as well as the transporter and trafficking protein Slc17a7 (Fold change -2.39), without significant differences compared to the DMSO-10 µM (**Figure 5A**). After treatment with benzothiazole **7g**, compared to DMSO-10 µM, the expression of Slc7a11 was significantly increased (Fold change is 4.39, p = 0.0036). It is important to note that Slc7a11 encodes a member of a heteromeric, sodium-independent, anionic amino acid transport system that is highly specific for cysteine and glutamate. Also, agent **7g** increased the expression of GABA receptors Gabrg1 and Gabrg3 and the transporter and trafficking protein Slc1a6 by more than 2-fold, while no significant difference was observed compared to the control group. Benzothiazole **7g** also decreased the trafficking protein Slc17a7 by more than 2-fold with no substantial differences compared to the DMSO-10 µM (**Figure 5B**). **Lipidomics analysis after benzothiazole treatment:** Lipidomics analysis provides in-depth insight into lipid mediators (LMs), a class of bioactive metabolites involved in many physiological processes.(*23*) Thus, the effect of the synthesized benzothiazoles on lipid profiling was investigated using a well-established and specific liquid chromatography/mass spectrometry (LC/MS) method.(*23, 24*)

**Figure 5.**
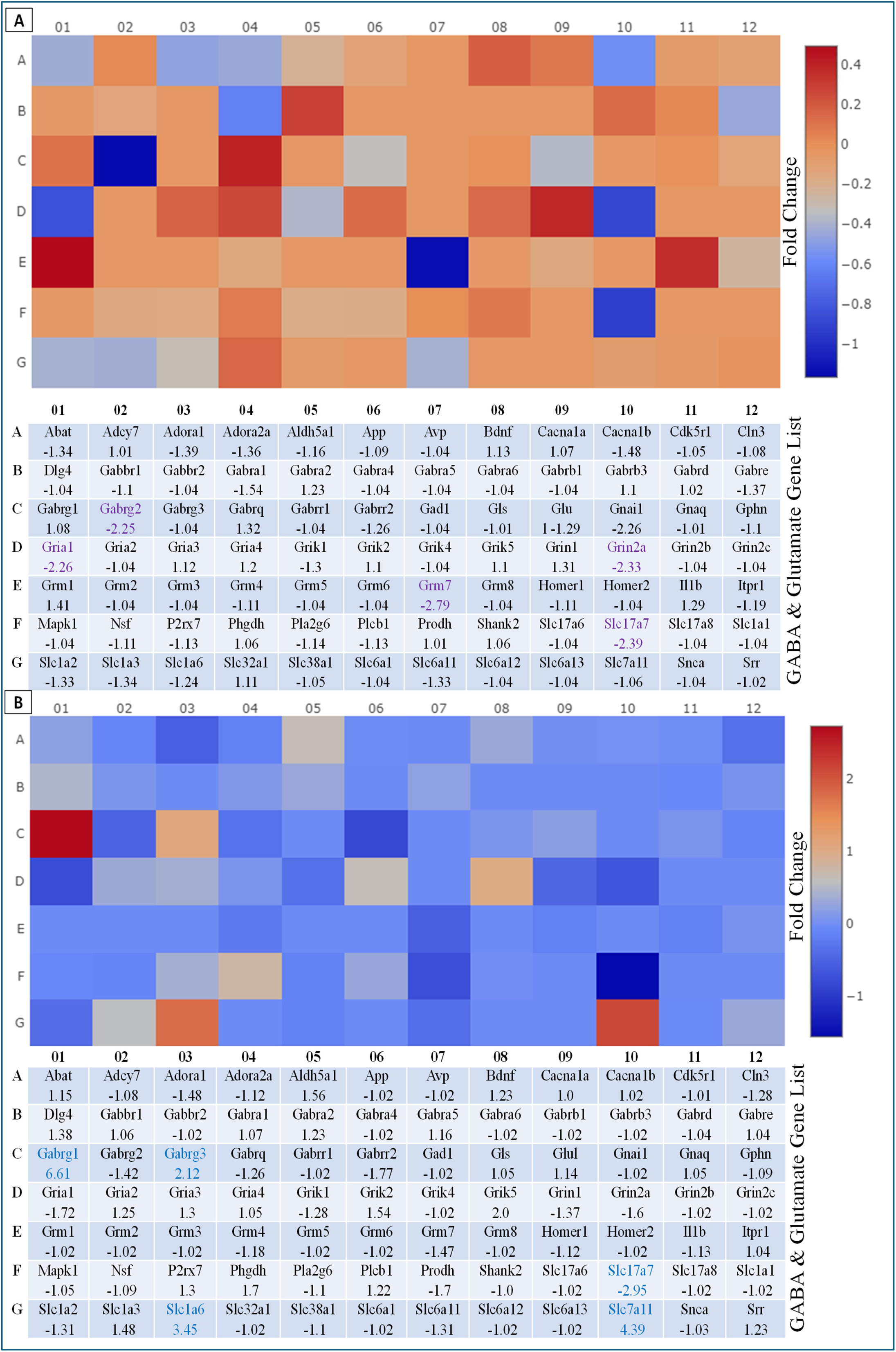
Detection of GABA receptors by Mouse GABA & Glutamate RT² Profiler PCR Array. A) GABA array for benzothiazole agent **7b** indicated more than 2-fold downregulation of Gabrg2, Grm7, Gria1, Grin2a, and Slc17a7. B) GABA array for benzothiazole agent **7g** indicated more than 2-fold upregulation of Gabrg1, Gabrg3, Slc7a11, Slc1a6, and downregulation of Slc7a7.

From the lipidomics quantification data shown in **Table 2**, which are then normalized to control values and presented as a heatmap in **S-Figure 1**, it is evident that agents **7b** and **7g** have a contrary effect on the lipid profile. Agent **7b** significantly decreased most quantified lipid mediators, while increasing only the linoleic acid (LA) pathway, especially 13-HODE and 9-HODE.(*69*) The 9-HODE and 13-HODE are lipid peroxidation products of linoleic acid oxidation/transformation into hydroperoxyl derivatives. These lipid peroxidation products suggest that agent **7b** could induce lipid peroxidation, which may explain the decreased number of healthy LMs. On the other hand, agent **7g** increased the total arachidonic acid (AA) pathway (PGE_2_, 6-keto-PGF1α, and 12-HHT), EPA, DHA, and only 9-HODE LM from the LA pathway. Benzothiazole **7g** increased the PGE_2_ concentration compared to the control and DMSO, while agent **7b** significantly decreased the PGE_2_ concentration compared to the control and DMSO groups (p = 0.007).

**Table 2:** Lipidomic analysis of the impact of synthesized benzothiazoles 7b and 7g on # cells.

| Quantified Analysis | LM | Concentration (Pg/mg protein) |  |  |  |
| --- | --- | --- | --- | --- | --- |
|  |  | Benzothiazole 7b | Benzothiazole 7g | DMSO | Control |
| Arachidonic acid (AA), n-6 polyunsaturated fatty acid (PUFA) | 6-keto-PGF1a | 572.11 | 2031.65 | 1002.12 | 1226.84 |
|  | PGF2a | 5.93 | 11.62 | 8.17 | 14.20 |
|  | PGE2 | 4.02 | 10.48 | 8.06 | 8.45 |
|  | PGD2 | 2.63 | 6.94 | 6.60 | 6.52 |
|  | 12-HHT | 276.20 | 1000.85 | 432.11 | 581.19 |
|  | AA | 21.57 | 101.36 | 48.88 | 100.90 |
|  | TOTAL | 882.46 | 3162.90 | 1505.95 | 1938.09 |
| Linoleic acid (LA), n-6 PUFA | 12,13-DIHOME | 4.98 | 4.54 | 4.33 | 4.57 |
|  | 9,10-DiHOME | 0.00 | 0.00 | 0.00 | 0.00 |
|  | 13-HODE | 3135.45 | 2760.36 | 1985.19 | 2535.38 |
|  | 9-HODE | 6059.07 | 6252.32 | 3691.36 | 4583.22 |
|  | TOTAL | 9199.50 | 9017.22 | 5680.88 | 7123.17 |
| Docosahexaenoic acid (DHA), n-3 PUFA | DHA | 634.29 | 1296.26 | 650.12 | 885.83 |
| Endocannabinoids and congeners | AEA | 0.00 | 0.00 | 0.00 | 0.00 |
|  | OEA | 29.69 | 43.80 | 38.07 | 62.98 |
|  | PEA | 58.30 | 74.72 | 53.00 | 86.99 |
|  | 2-AG | 65.95 | 112.12 | 110.42 | 209.62 |
|  | TOTAL | 153.94 | 230.63 | 201.49 | 359.58 |
| Eicosapentaenoic acid | EPA | 316.15 | 1021.14 | 565.11 | 835.02 |
|  | AA/EPA | 0.07 | 0.10 | 0.09 | 0.12 |

#### Intracellular ROS under oxidative stress conditions

The level of intracellular ROS in C2C12 cells was quantified to study the effect of benzothiazoles on ROS levels and investigate their potential antioxidant activity. A monolayer of C2C12 cells was pre-treated or co-treated with 10 µM of agents **7b** and **7g**, then 100 µM of H_2_O_2_ was used to induce free oxygen species. Cells treated with 100 µM of H_2_O_2_ only were used as a negative control, while normal cells were used as a positive control. The intracellular ROS were visualized using a carboxy-H2DCFDA fluorescence probe that produces a fluorescence signal upon interaction with ROS in the cells, as shown in **Figure 6A**. The fluorescence images were quantified using ImageJ software and presented as the ROS fold change relative to the control. Fluorescent images of the control indicated minimal to no ROS activity inside the cells, while the negative control (H_2_O_2_-treated cells) showed an increased fluorescent signal. The quantification indicated a significant increase in the intracellular ROS level, with a 2.5-fold increase compared to the positive control (p <0.0001). On the other hand, cells treated with **7b** and **7g** benzothiazole agents showed a significant decrease (p < 0.001) in intracellular ROS levels compared to the negative control (H_2_O_2_-treated cells), as shown in **Figure 6B**. It is important to note that benzothiazole groups completely quenched the effect of H_2_O_2_ and scavenged the produced ROS, and no significant difference was observed compared to the positive control. The quantitative analysis also revealed that agent **7b** acts as a scavenger when used as a pre- or co-treatment. In contrast, agent **7g** showed a better effect when used as a pre-treatment.

**Figure 6:**
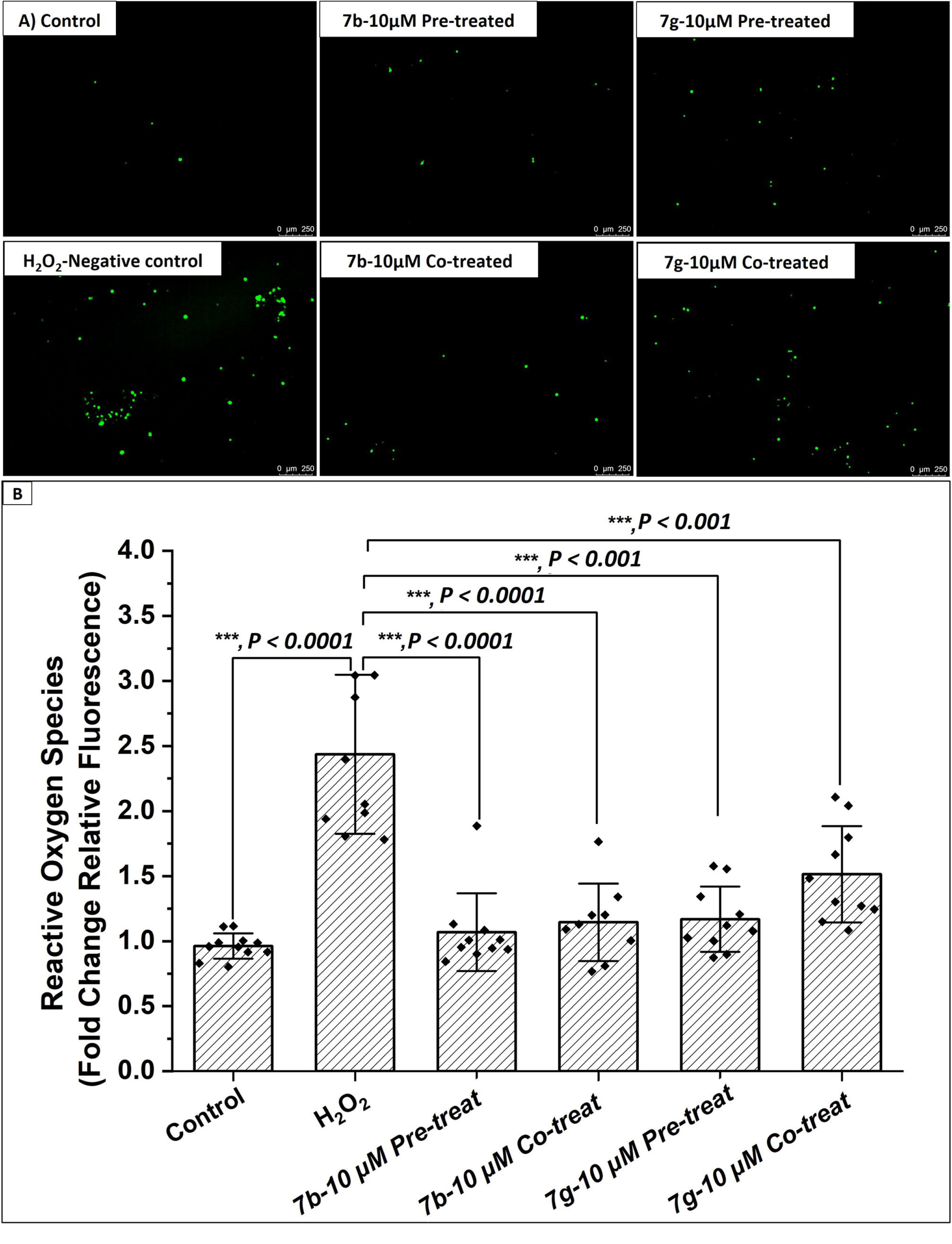
Benzothiazole agents 7b and 7g significantly decreased the level of intracellular ROS under oxidative stress conditions. (A) Fluorescent microscopy images visualize the intracellular ROS identified by the reaction with the probe carboxy-H2DCFDA. (B) Quantification of intracellular ROS indicated that the H_2_O_2_ treatment significantly increases the ROS level in the C2C12 cells within 30 minutes, while pre-treatment and co-treatment with benzothiazole agents 7b and 7g significantly mitigate the ROS level with potential antioxidant activity. One-way ANOVA with Tukey’s post-hoc test was performed for multiple comparisons between groups (* p<0.05, ** p<0.01, and *** p<0.001).

### 3.3 Molecular Docking

Molecular modeling is an indispensable step in modern drug discovery and design, especially structure-based molecular docking. The technique significantly contributes to identifying new drug candidates and understanding their structure-activity relationship. Here, the docking study helps us interpret the observed activity of the synthesized benzothiazoles and their effects on the Gabrg2 gene. Acetylated benzothiazole 7b significantly downregulated the Gabrg2 gene compared to benzothiazole 7g, providing a new perspective on a potential class of molecules that might treat epilepsy. Thus, the presented molecular docking study uses the molecular operating environment (MOE 2019.01) to visualize the binding interactions of these two compounds (7b and 7g) and their parent benzothiazole 7a. The binding scores and interactions are analyzed and compared with those of the crystalized anti-epileptic drug bound to the protein (PDB: 6X3X). The top-ten poses of benzothiazole conformations are analyzed to visualize how the acetyl group might affect binding in the pocket site of the targeted protein. Docking of the benzodiazepine drug with 6X3X protein(70) revealed the importance of the primary amino acids in the pocket site. Thus, three out of ten drug conformers have been found to interact with one or both amino acids (Thr C262, Asn 265, and Pro D233, Supplementary materials Table 1, Entry 1-3). **Figure 7** reveals the binding interactions of two benzodiazepine conformers with the amino acids in the active site, showing the possible interaction with Pro 233 via Pi-H interactions, where the aromatic π-electron system from a ligand interacts with the hydrogen atom of Pro 233, and Asn 265 via hydrogen bond (HB) interactions (**Figure 7a**, binding interactions S = - 6.0381 kcal/mol) and with Thr 262 via HB interactions (**Figure 7b**, binding interactions S = - 6.3367 kcal/mol). Interestingly, all docked benzothiazole derivatives show similar binding affinity compared to the benzodiazepine drug. However, not all exhibit similar interactions with the targeted amino acids. The parent benzothiazole (7a) docks with Pro 233 and Thr 262 via Pi-H interactions, exhibiting higher binding affinity than benzodiazepine (Supplementary materials **Table 1**, Entries 4-5). Meanwhile, the HB interaction of 7a with Asn 265 shows a similar binding affinity to that of the reference compound (**Figure 7c**, S = - 6.3613 kcal/mol, Supplementary materials Table 1, Entry 6). Similar observations were recorded upon docking 7g conformers (**Figure 7d**, S = - 6.6646 kcal/mol, Supplementary materials Table 1, Entry 7-9). Interestingly, Pi-H interactions of thiazole rings in all the benzothiazole 7b conformers result in better binding affinity compared to hydrogen bond interactions with the targeted amino acids (**Figure 7e**, S = -7.715 kcal/mol; Supplementary materials Table 1, Entry 10-14). Acetylated compounds belong to a fascinating class of prodrugs that enhance pharmacokinetics, stability, and chemical properties, particularly in the central nervous system. In-vivo or in-vitro hydrolysis of benzothiazole 7b may result in a 4-hydroxy benzothiazole derivative (**Figure 7f**). Therefore, a docking study of the free hydroxy benzothiazole was performed to confirm the similarity of binding interactions within the active site and to explore the potential differences between the free hydroxy and acetylated masked benzothiazole. Remarkably, the free hydroxy benzothiazole retains binding to the targeted amino acid Pro 233 via thiazole interaction and demasking the acetylated group results in an alternative hydrogen-bond interaction with Thr 262 (**Figure 7f**, S = - 6.3367 kcal/mol, Supplementary materials Table 1, Entry 15).

**Figure 7.**
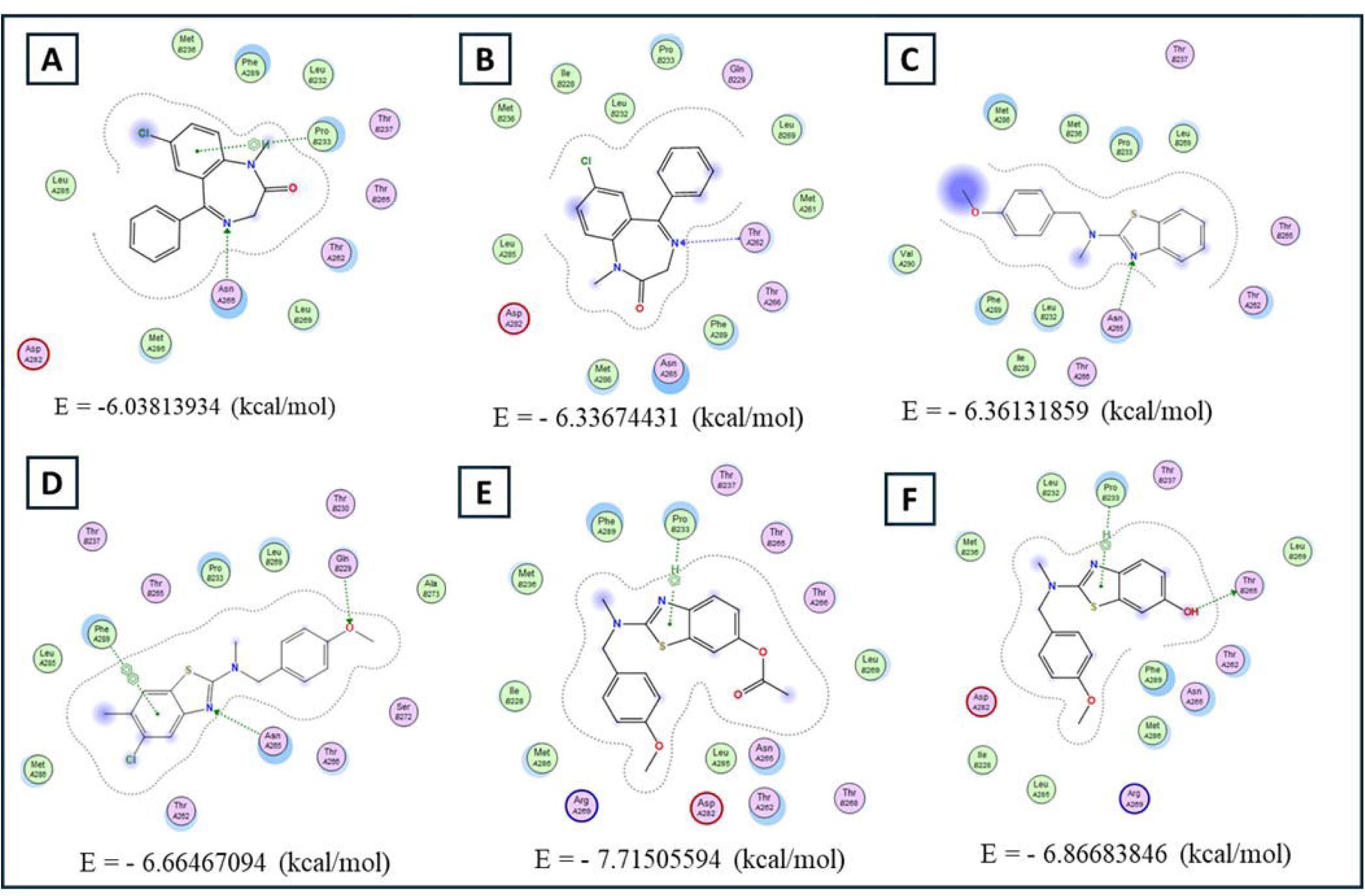
Docking of anti-epileptic drug (benzodiazepine) and benzothiazoles with co-crystallized protein (PDB: 6X3X) using MOE software. (A and B) 2D Molecular docking of benzodiazepine conformers. (C) 2D Molecular docking of benzothiazole **7a**. (D) 2D Molecular docking of benzothiazole **7g**. (E) 2D Molecular docking of acetylated benzothiazole **7b**. (F) 2D Molecular docking of 4-hydroxy-benzothiazole

### 3.4 Overall Proposed Pathway Analysis

Using the Qiagen Ingenuity Pathway Analysis (IPA) software to integrate lipidomics and gene array data, we constructed the proposed pathway shown in **Figure 8**. Benzothiazole **7b** significantly downregulates genes such as GRIA1, GRIN2A, GRM7, and SLC17A7, which are linked to glutamatergic and GABAergic synaptic signaling in skeletal muscle. These changes result in altered neurotransmitter profiles, as highlighted by reduced GABA and changes in aminobutyric acid concentrations. Downregulation of these neurotransmitter-linked genes intersects with key metabolic and lipid mediators, most notably PGE_2_. In the illustrated network, PGE_2_ integrates signals from cytokines (TNF, IL-1β), kinases (MAPK, ERK), and eicosanoid metabolism (via PTGS2 and PLA2), ultimately driving the expression of genes that support skeletal muscle myogenesis (e.g., VEGFA, VIM, SP1).

**Figure 8:**
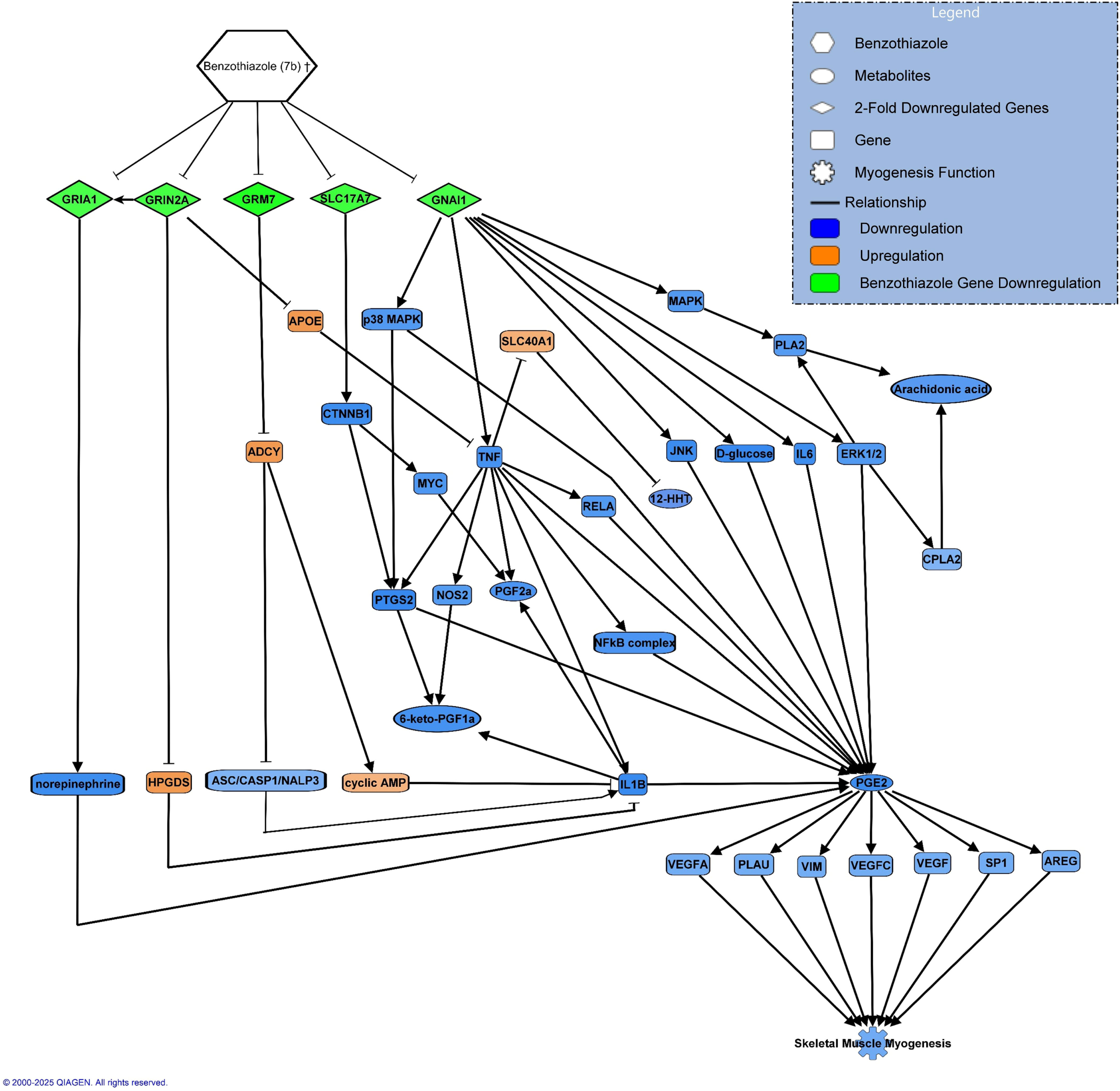
Benzothiazole derivatives, particularly agents **7b** and **7g**, exert significant and divergent regulatory effects on skeletal muscle myogenesis, as demonstrated by integrative analysis of gene expression, lipidomics, and functional myogenesis assays. The pathway map illustrates the upstream impact of benzothiazole (**7b**) on key neurotransmitter and metabolic genes, as well as its downstream modulation of prostaglandin (PGE_2_) signalling, a central key in myogenic regulation.

## 4. DISCUSSION

Our synthetic findings collectively indicate that the newly developed hypervalent iodine–mediated cyclization offers a concise and environmentally benign route to benzothiazole scaffolds from simple thioureas, in contrast to classical methods that rely on harsher conditions and longer reaction times. The dependence of chemoselective acetylated benzothiazole formation on IBDA equivalents underscores a tunable system in which 2.0 equivalents favor acetylated products (**7b** and **7d**), while 1.0 equivalents afford non-acetylated benzothiazoles (**7a** and **7c**) in good to moderate yields. This flexibility in product outcome is particularly relevant for structure–activity relationship (SAR) exploration, as exemplified by the distinctive biological profiles of **7b** and **7g**.

The cytotoxicity and myogenesis data together suggest that benzothiazole derivatives can be applied safely at low micromolar concentrations (5–10 µM) in skeletal muscle models, where they modulate differentiation rather than induce cell death. The absence of cytotoxicity at these doses, combined with the enhanced proliferation seen for **7b**, **7c**, **7e**, and **7g**, positions these scaffolds as a tractable platform for pro-myogenic or anti-myogenic agents depending on substitution. The robust suppression of the fusion index by **7b** at both 5 and 10 µM, in contrast to the neutral profile of **7g** at the same doses, reveals early evidence of a divergence in myogenic regulatory potential rooted in chemical structure.

The metabolomics data further support the concept that **7b** and **7g** exert distinct effects on the GABA–BAIBA axis in C2C12 cells. For **7b**, the pronounced reduction in GABA combined with the selective increase in L-BAIBA in both cells and conditioned media suggests a shift away from inhibitory neurotransmitter abundance toward a myokine-like metabolite associated with mitochondrial biogenesis and metabolic adaptation. The absence of major changes in D-BAIBA and the more modest effect of **7g** on GABA underscore that **7b** uniquely reprograms aminobutyric acid metabolism. Such a profile is consistent with an anti-excitatory, anti-inflammatory signature that may influence both myogenic and neuronal phenotypes.

To investigate these metabolomic observations further, we performed a gene array. This array presents genes essential for synthesizing and transporting GABA, glutamate, and downstream signaling. The gene list of this array includes 17 GABAergic Synapse “Neurotransmitter Receptors,” 9 Signaling Downstream of GABAergic Synapse, 20 Glutamatergic Synapse, 15 Signaling Downstream of Glutamatergic Synapse, 16 Transporters/Trafficking Proteins, and 7 metabolism genes. The data reinforce these metabolic observations by showing that **7b** downregulates Gabrg2 and key glutamatergic receptors and transporters (Gria1, Grin2a, Grm7, Slc17a7), supporting suppression of both GABAergic and glutamatergic signaling pathways at the receptor and transporter level. This broad downregulation may contribute to decreased synaptic excitability and supports the concept of **7b** as a potential modulator of epileptiform activity, notwithstanding the cost of reduced myogenic signaling in muscle. By contrast, **7g** upregulates Slc7a11, Gabrg1, Gabrg3, and Slc1a6, indicating a different regulatory axis that enhances cystine/glutamate transport and GABA receptor expression without significantly altering control-level expression patterns. Thus, the PCR array results suggest that agent **7b** significantly regulates GABA receptor Gabrg2, while agent 7g is a potential target for regulating Slc7a11. Slc7a11 is a key cystine/glutamate transporter that regulates cellular lipid peroxidation and restrains ferroptosis.(*70*) The selective induction of Slc7a11 by **7g** is particularly significant, given this transporter’s established role in cystine uptake, glutathione synthesis, and ferroptosis resistance, thereby linking **7g** to antioxidant and cytoprotective mechanisms. These findings highlight the therapeutic potential of benzothiazoles in targeting both GABAergic and glutamatergic pathways, supporting their development for genetic or non-genetic epilepsy and CNS disorders, involving synaptic dysfunction or ferroptosis mechanisms.

Lipid mediators are directly associated with inflammation, oxidative stress, tissue healing, and regeneration, and can regulate the mass and function of skeletal muscles overall.(*26, 71*) LMs derived from polyunsaturated fatty acids (PUFAs) can be classified based on their structure into omega-3 (ω-3) PUFAs, which include Eicosapentaenoic Acid (EPA), docosahexaenoic acid (DHA), and α-linoleic acid (ALA), and omega-6 (ω-6) PUFAs that contain the arachidonic acid (AA) pathway. It is reported that ω-3 and ω-6 PUFAs affect the body’s metabolic functions differently. ω-6 PUFAs are generally associated with inflammatory responses, constriction of blood vessels, and platelet aggregation.(*72, 73*) On the other hand, ω-3 PUFAs have anti-inflammatory and pro-resolving activities.(*72, 74*) Quantifying these LMs can reveal their effect and possible signaling pathways under benzothiazole treatment conditions. Our lipidomics findings highlight that **7b** and **7g** exert opposite effects on eicosanoid and ω-3/ω-6 lipid mediator networks. The **7b** agent generally depletes multiple lipid mediators while increasing 9-HODE and 13-HODE, consistent with elevated lipid peroxidation and reduced “healthy” resolution-phase lipid mediators. In contrast, 7g elevates AA-derived prostanoids (PGE_2_, 6-keto-PGF1α, 12-HHT) as well as EPA and DHA, suggesting an environment conducive to pro-regenerative, pro-resolving signaling.

Most importantly, Prostaglandin E2 (PGE_2_) is a major PG produced from AA via the cyclooxygenase 1/2 (COX1/2) pathways. It accelerates skeletal muscle myogenic differentiation and regeneration by promoting proliferation and increasing intracellular ROS production in myoblasts.(*24, 26–28, 73, 75*) The 6-keto prostaglandin F1α (6-keto-PGF1α) is the inactive, non-enzymatic hydrolysis product of prostacyclin or prostaglandin I2 (PGI2), which is an effective vasodilator due to its action in the prevention of platelet aggregation.(*75*) 12-hydroxyheptadecatrienoic acid (12-HHT) is a ligand for the leukotriene B4 receptor type 2 (BLT_2_) and has recently been reported to play a significant role in wound healing.(*76*) Liu et al. reported that 12-HHT/BLT_2_ induces keratinocyte migration and wound closure through NF-κB-dependent up-regulation of TNF and matrix metalloproteinase 9 (MMP9).(*76*) Of particular importance, PGE_2_ emerges as a central bioactive lipid modulated in opposite directions by agents **7b** and **7g**. Recent studies have elucidated that transient surges in PGE_2_ following injury act directly on muscle stem cells (MuSCs) via the EP4 receptor to drive robust proliferation and accelerate muscle repair and strength recovery.(*24*) Loss of PGE_2_ signaling, including pharmacologic NSAID inhibition, impairs MuSC expansion, reduces muscle force, and compromises regenerative capacity.(*26*) A similar effect was induced by agent **7b**, indicating downregulation of PGE_2_. Conversely, augmentation of PGE2-induced biosynthesis potentiates MuSC activation and myogenesis, with increased muscle mass and strength demonstrated in animal models(*26–28, 42*). The dramatic modulation of PGE_2_ by the two agents directly aligns with the observed phenotype. Enhanced myogenic differentiation and slightly elevated ROS levels in cells treated with agent **7g** are tightly linked to elevated PGE2, promoting proliferation and downstream myogenic differentiation. In contrast, the dramatic reduction in myogenic differentiation after **7b** is explained by a significant reduction in intracellular PGE2, thereby impairing myogenic signaling. The present data thus reinforces the paradigm in which benzothiazole derivatives serve as molecular switches at the intersection of lipid signaling and muscle biology, with PGE_2_ acting as the pivotal factor for myogenic outcomes.

Through overall comparison of agents **7b** and **7g**, we can observe that agent **7b** exerted a more prominent inhibitory effect on myogenesis compared to **7g**, likely by blunting key inflammatory signaling pathways that initiate muscle differentiation. This agent specifically reduced GABA levels while elevating BAIBA, reflecting its anti-inflammatory profile. Lower GABA contributes to impaired myogenic signaling, while elevated BAIBA—derived from enhanced ω-3 influences—activates mitochondrial biogenesis via PPAR-α, incorporating into mitochondrial membranes to improve electron transport chain efficiency and ATP production. Agent **7g** increased both DHA and EPA, which are known for their beneficial effects on skeletal muscle. EPA and DHA were reported to prevent the development of cardiovascular diseases, lipotoxicity, inflammation, and metabolic abnormalities in skeletal muscle.(*77, 78*) Further, as previously reported, DHA has a specific capacity to modulate redox mechanisms and reduce lipid peroxidation.(*79*) In particular, the increase in DHA lipid mediator levels with agent **7g** was paralleled by overexpression of the Slc7a11 gene, suggesting synergistic regulation of glutathione synthesis and antioxidant defense, highlighting the potential of agent **7g** not only to promote myogenesis via PGE_2_ signaling but also to protect skeletal muscle via enhanced redox homeostasis and mitigation of lipid peroxidation.

Furthermore, the ROS assays indicate that both **7b** and **7g** possess potent antioxidant capacity, fully quenching H2O2-induced ROS to levels comparable to untreated controls. For **7b**, this antioxidant behavior coexists with suppressed PGE2 and extensive downregulation of myogenesis- and neurotransmission-related genes, suggesting a profile that combines anti-inflammatory, anti-oxidative, and anti-myogenic effects. For **7g**, ROS mitigation is accompanied by increased PGE2, elevated EPA/DHA, and upregulation of Slc7a11, pointing to a complementary profile that supports both redox homeostasis and regenerative signaling. The preferential efficacy of **7g** as a pre-treatment in ROS assays further underscores its potential utility in prophylactic or conditioning paradigms where oxidative insults are anticipated.

The docking studies provide a structural rationale for the observed transcriptional and functional effects on Gabrg2 and related targets. **7b** exhibits stronger binding affinities than both **7a**, **7g**, and the reference benzodiazepine in the 6X3X binding pocket, relying heavily on Pi–H interactions through the thiazole ring with key residues such as Pro 233, Thr 262, and Asn 265. The absence of HB interactions in **7b’s** bound poses suggests that π-stacking and hydrophobic contacts dominate its binding mode, consistent with potent but potentially non-classical modulation of receptor conformation. Hydrolysis of **7b** to the free hydroxy analogue maintains interactions with Pro 233 while introducing a new HB to Thr 262, implying that both the acetylated prodrug and its deprotected form can occupy the pocket in complementary ways. These docking outcomes align with the marked downregulation of Gabrg2 expression in cells treated with **7b** and support classifying this compound as a promising Gabrg2-targeting scaffold. The integrated pathway analysis suggests that the combined alterations in gene expression, metabolite profiles, and lipid mediators converge on a PGE2-centered network that coordinates myogenesis, inflammatory signaling, and neurotransmission in skeletal muscle. **7b**-driven downregulation of GRIA1, GRIN2A, GRM7, and SLC17A7, together with reduced PGE2 and other prostanoids, positions 7b as a suppressor of myogenic and synaptic signaling networks. In contrast, **7g** enhances PGE_2_ and upregulates genes such as Gabrg1, Gabrg3, Slc7a11, and Slc1a6, thereby promoting a milieu supportive of regeneration, antioxidant defense, and altered GABAergic signaling. The dual actions of these agents—as both myogenic and synaptic modulators—highlight benzothiazoles as versatile tools for dissecting cross-talk between skeletal muscle and neurotransmitter systems.

Taken together, our data show that benzothiazole agents act as dual regulators of myogenic differentiation via PGE_2_-centered and neurotransmitter signaling pathways, affecting both gene expression and the lipid environment in skeletal muscle. Lipidomic analyses support pathway predictions: treatment with benzothiazole **7b** leads to notable reductions in key lipid mediators (e.g., 6-keto-PGF1α, PGF2α, PGE_2_), while **7g** increases these bioactive lipids, reflecting opposite impacts on the lipid signaling environment of muscle cells. This is particularly relevant for PGE_2_, whose abundance is tightly linked to myogenic progression and is coordinated through the regulatory networks depicted in the pathway figure. Further, gene array data reveal pronounced (>2-fold) downregulation of myogenesis-linked and neurotransmission genes following **7b** exposure (GRIA1, GRIN2A, GRM7, SLC17A7), in line with the pathway diagram’s green highlights. This genetic repression is associated with anti-inflammatory effects, as benzothiazoles **7b** and **7g** mitigate hydrogen peroxide-induced ROS in myotubes, demonstrating antioxidant action alongside their impact on differentiation and signaling.

The contrasting biological effects of **7b** and **7g** on PGE_2_ and GABA arise from their distinct chemical structure and substitution patterns. We speculate that the acetate and methoxy groups of **7b** inhibit bioactive lipid synthesis and gene expression, while the chloro and methyl groups of **7g** enhance binding to biosynthetic and signaling proteins, increasing PGE_2_ production and promoting myogenic progression. The acetate substitution in **7b**, as a polar functional group, can increase hydrophilicity and may introduce steric effects that interfere with the site-specific binding of enzymes or receptors.(*80–82*) Thus, we believe that the presence of an acetate group (as in **7b**) can introduce steric and electronic constraints, reducing substrate or inhibitor access to enzyme active sites (such as COX) and thereby diminishing PGE_2_ synthesis. On the other hand, in agent **7g**, the chloro group is electron-withdrawing, while the methyl is electron-donating. This combination can stabilize enzyme–ligand interactions or modulate binding to active sites, including cyclooxygenase (COX) and other lipid/metabolic enzymes. A recent study shows that chloro- and methyl-substitutions at C6 of benzothiazoles enhance biological activity.(*83*) Furthermore, other studies have reported that electron-withdrawing substituents enhance the inhibitory activity or binding of COX-2 inhibitors by improving the inhibitor’s stabilization and orientation in the enzyme’s active site.(*84, 85*) Electron-donating groups (like methyl) provide favorable hydrophobic contacts and help tune the compound’s polarity/stability within the binding pocket. Thus, we speculate that agent **7g** favors binding to myogenic or lipid biosynthetic machinery, thereby augmenting PGE_2_ production and transcription of genes associated with regeneration (Gabrg1, Gabrg3, Slc7a11, Slc1a6).

Overall, this study reveals new functional diversity within the benzothiazole scaffold, demonstrating precise structure-activity relationships that regulate GABAergic neurotransmission, lipid mediator signaling, and skeletal myogenesis. The acetylated derivative **7b** emerged as a robust suppressor of myogenic gene networks and prostaglandin E2 synthesis, coupled with GABA downregulation and unique modulation of L-BAIBA production. Conversely, the 5-chloro-6-methyl derivative **7g** promoted myogenic differentiation, increased PGE_2_, and upregulated genes related to antioxidants and neurotransmitters, with corresponding elevations in omega-3 and omega-6 lipid mediators. Both compounds exhibited pronounced antioxidant capacities in cellular assays. Pathway integration established that coordinated modulation of PGE_2_ and GABAergic signaling underpins the dual myogenic and synaptic actions of these molecules. Taken together, these results establish benzothiazole derivatives as promising molecular tools to dissect and potentially manipulate the biochemical crossroads of skeletal myogenesis and neurotransmitter balance, laying the foundation for targeted therapeutic strategies in muscle and synaptic disorders. However, we investigated a small benzothiazole library; the outstanding results encourage further exploration of various benzothiazole structures in the near future.

## Conclusion

Our thorough in vitro assessment showed that benzothiazole derivatives, at low micromolar concentrations, are non-cytotoxic and have different effects on myogenic differentiation. Agent **7b** strongly inhibited myotube formation and decreased key genes in both myogenic and neurotransmitter pathways, notably lowering PGE2, GABA, and synaptic gene expression, while also increasing antioxidant and anti-inflammatory responses. In contrast, agent **7g** promoted myogenesis, elevated PGE2 and pro-regenerative lipid mediators, and upregulated genes related to antioxidant defense, neurotransmitter transport, and regenerative ability. Metabolomic, lipidomic, and gene array analyses collectively confirm that the interaction between PGE2 and GABAergic signaling forms a central regulatory network influenced by these compounds. The unique structural features of **7b** and **7g** account for their distinct effects on lipid mediator profiles, neurotransmitter levels, and gene networks, providing insight into how specific benzothiazole modifications lead to distinct biological outcomes.

Both agents also showed strong antioxidant activity by neutralizing ROS in muscle cells, emphasizing their potential to protect against oxidative stress. Molecular docking studies further explained the structural basis for the observed gene-regulatory effects, especially the high binding affinity of **7b** for Gabrg2 and related targets. Overall, this work suggests that benzothiazole derivatives are useful chemical tools for exploring biochemical communication between muscle and synaptic biology. Their dual action, guided by detailed structure–activity relationships, opens possibilities for future therapeutic development targeting muscle and neurological disorders. More research and optimization of the benzothiazole scaffold are needed to fully realize its biomedical potential.

## Acknowledgments

## Funding

The authors thank the following for generous financial support

National Science Foundation grant CH (CJL)

The Robert A. Welch Foundation grant Y-1362 (CJL)

National Institutes of Health grant R01DE031872

National Institutes of Health grant S10OD025230)

National Institutes of Aging 5R01AG056504-02

National Institutes of Aging 1R56AG049083-01

National Institutes of Aging 5P01AG039355-08)

Trauma Research and Combat Casualty Care Collaborative (TRC4) by the UT System

The George W. and Hazel M. Jay Research Endowments

UTSW-NCATs grant # 1UL1TR003163-04

The authors are grateful to the UTA Shimadzu Institute for Research Technologies and to the Bone-Muscle Research Center Core Facilities at UTA.

## Author contributions

Conceptualization: MA, KA, MB, CJL

Methodology: MA, KA, JH, ZW

Investigation: MA, KA, JH, ZW

Visualization: MA, KA, JH, ZW, VV

Supervision: VV, CJL, MB

Writing—original draft: MA, KA, JH, ZW

Writing—review & editing: MA, KA, JH, ZW, VV, CJL, MB

## Data and materials availability

All data are available in the main text or the supplementary materials

