## Supplemental S-Figure 1 for "Benzothiazole Derivatives as Dual Modulators of PGE2 and GABAergic Signaling in Skeletal Muscle"

Marian N. Aziz *et al.*

**This file includes:**

**Figs. S-Figure 1**

**Tables: S-Table 1**

**Spectra**: **Data S3 to S14**

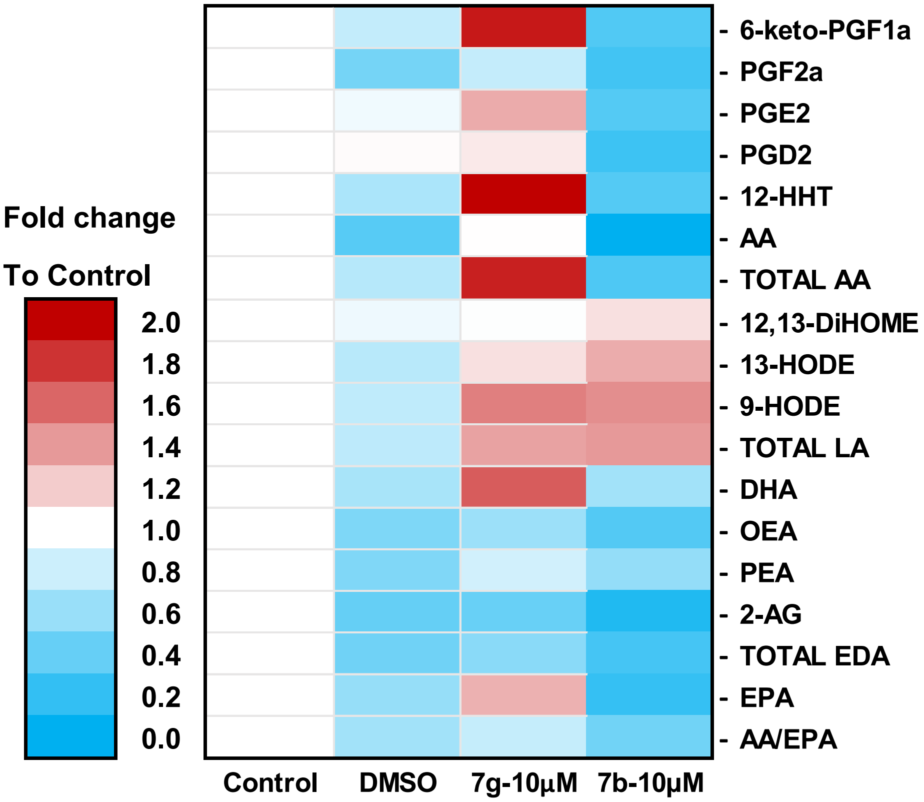


**S-Figure 1: Benzothiazole agents 7b and 7g regulate the lipid profile in C2C12 skeletal muscle cells.** Heatmap of quantified LMs normalized to positive control indicate that benzothiazole agent **7b** significantly decrease the lipid profile while agent **7g** increase the LMs concentration compared to other groups.

### Supplementary materials Table 1

| | 2-D docking poses | Binding affinity | Interactions Details | | | | |
|---|---|---|---|---|---|---|---|
| | | | Ligand | Receptor | Interactions | Distance | E (kcal/mol) |
| 1 | 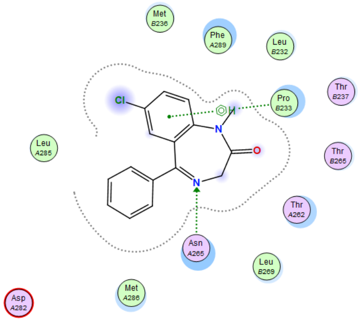 | -6.03813934 | N16 | ASN 265 | H-acceptor | 2.84 | -2.1 |
| | | | 6-ring | PRO 233 | pi-H | 3.75 | -0.6 |
| 2 | 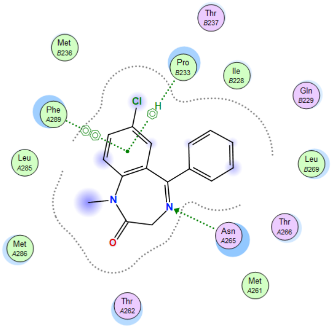 | -6.42177391 | 6-ring | PRO 233 | pi-H | 4.14 | -0.5 |
| | | | 6-ring | PHE 289 | pi-pi | 3.77 | -0.0 |
| 3 | 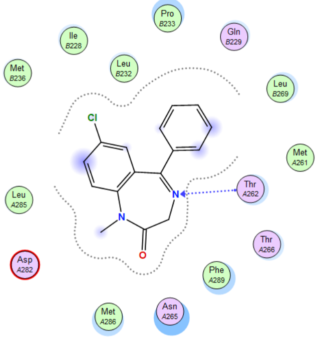 | -6.33674431 | N16 | THR 262 | H-acceptor | 3.49 | -0.6 |
| 4 | 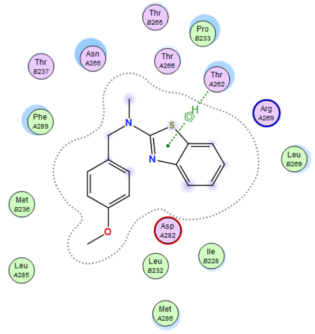 | -6.9812212 | 5-ring | THR 262 | pi-H | 3.64 | -0.5 |

| | | | | | | | |
|---|---|---|---|---|---|---|---|
| **5** | 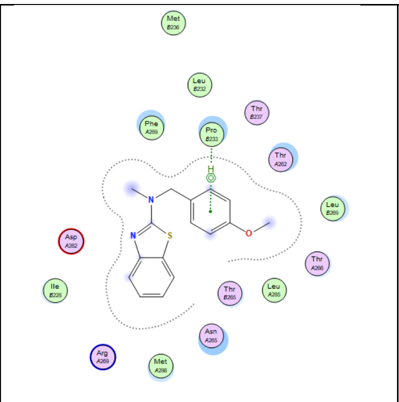 | -6.74971676 | 6-ring | PRO 233 | pi-H | 3.77 | -0.6 |
| **6** | 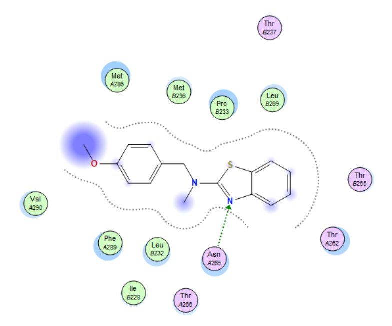 | -6.36131859 | N17 | ASN 265 | H-acceptor | 3.23 | -0.5 |
| **7** | 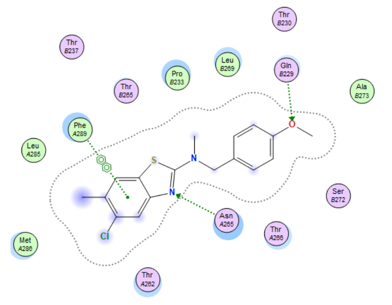 | -6.66467094 | N17 | ASN 265 | H-acceptor | 3.47 | -0.8 |
| | | | O31 | GLN 229 | H-acceptor | 2.81 | -0.7 |
| | | | 6-ring | 6-ring/PHE 289 | pi-pi | 3.91 | -0.0 |
| **8** | 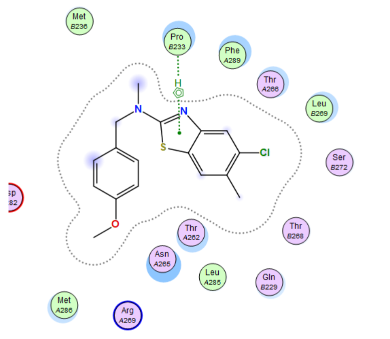 | -6.72870636 | 5-ring | PRO 233 | pi-H | 3.75 | -0.6 |

| | | | | | | | |
|---|---|---|---|---|---|---|---|
| **9** | 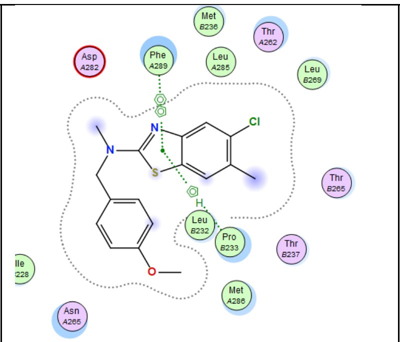 | -6.72259903 | 5-ring | PRO  233 | pi-H | 3.82 | -0.8 |
| | | | 5-ring | 6-ring/PHE 289 | pi-pi | 3.98 | -0.0 |
| **10** | 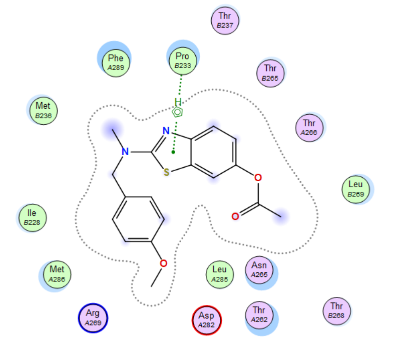 | -7.71505594 | 5-ring | PRO  233 | pi-H | 3.73 | -0.7 |
| **11** | 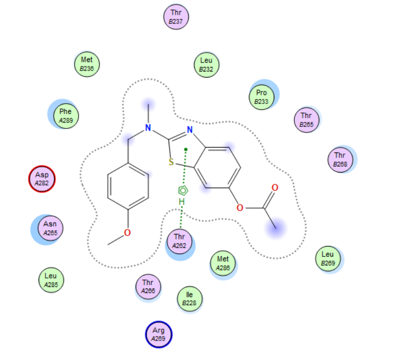 | -7.6685605 | 5-ring | THR  262 | pi-H | 3.96 | -0.5 |
| **12** | 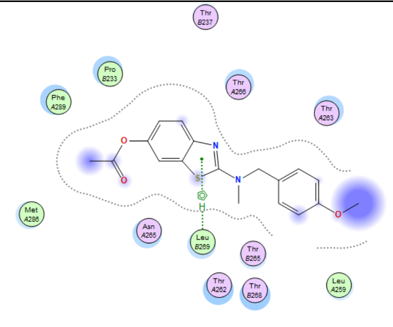 | -7.6685605 | 5-ring | LEU  269 | pi-H | 3.88 | -0.5 |

| | | | | | | | |
|---|---|---|---|---|---|---|---|
| **13** | 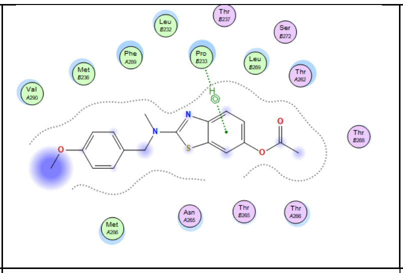 | -7.01082182 | 6-ring | PRO 233 | pi-H | 4.17 | -0.5 |
| **14** | 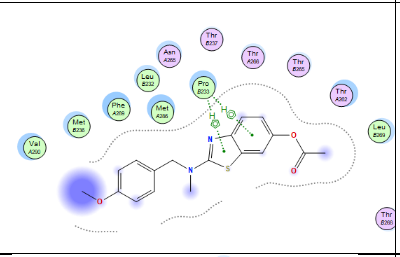 | -6.92033482 | 6-ring | PRO 233 | pi-H | 3.74 | -0.5 |
| | | | 5-ring | PRO 233 | pi-H | 4.05 | -0.6 |
| **15** | 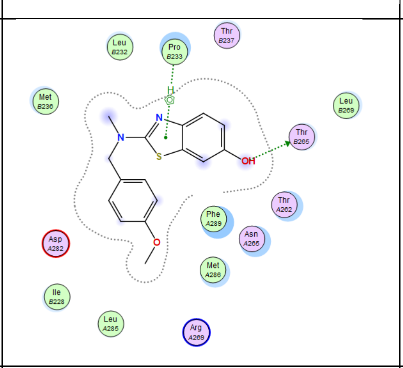 | -6.86683846 | O27 | THR 265 | H-donor | 3.07 | -0.7 |
| | | | 5-ring | PRO 233 | pi-H | 3.74 | -0.9 |
| **16** | 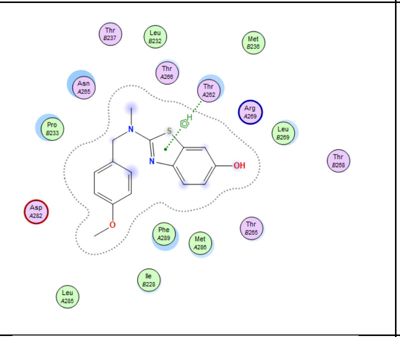 | -6.97476816 | 5-ring | THR 262 | pi-H | 3.61 | -0.5 |
| **18** | 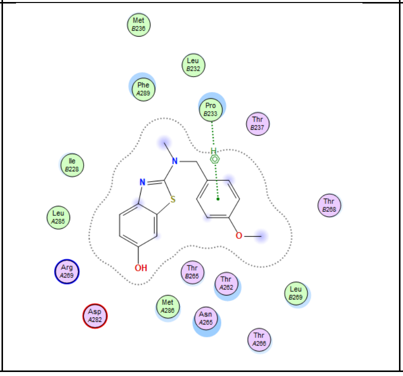 | -6.77980042 | 6-ring | PRO 233 | pi-H | 4.08 | -0.7 |

**Supplementary materials Data S3-S14: $^{1}$H and $^{13}$C NMR spectra for:**

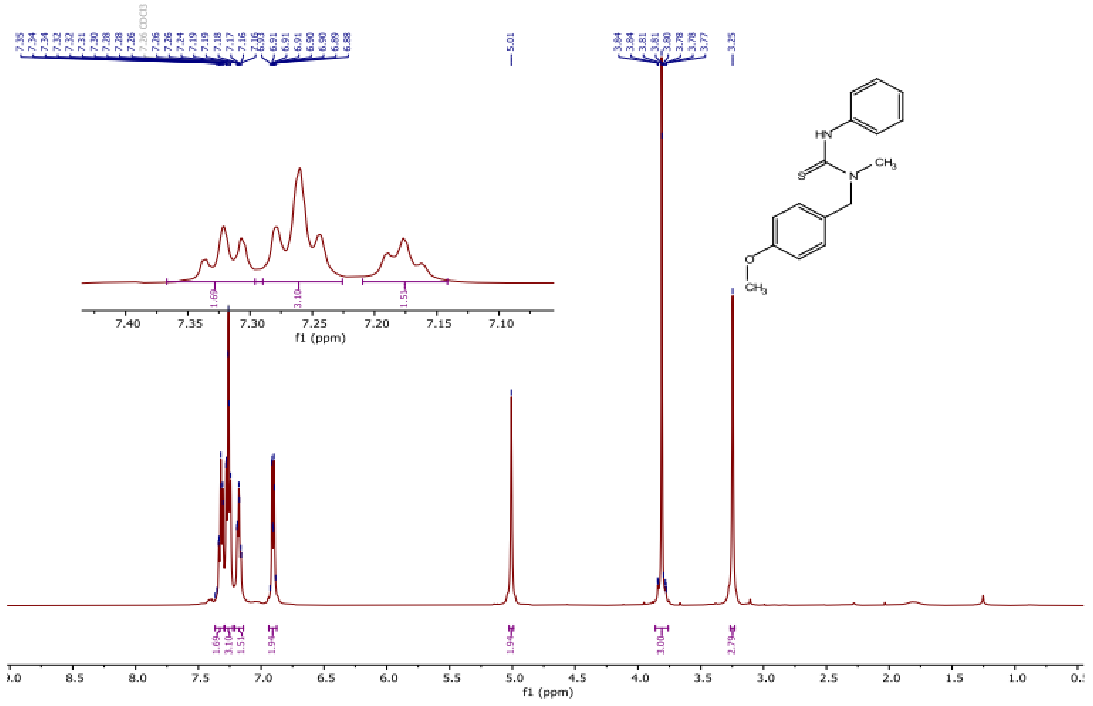


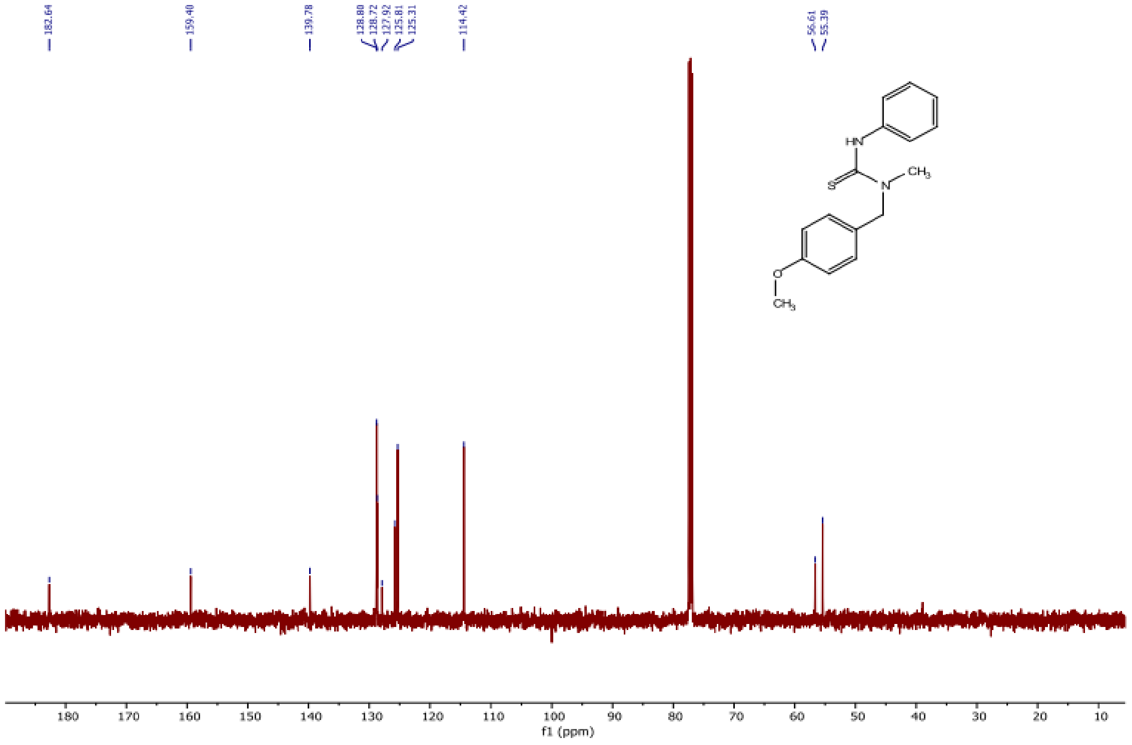

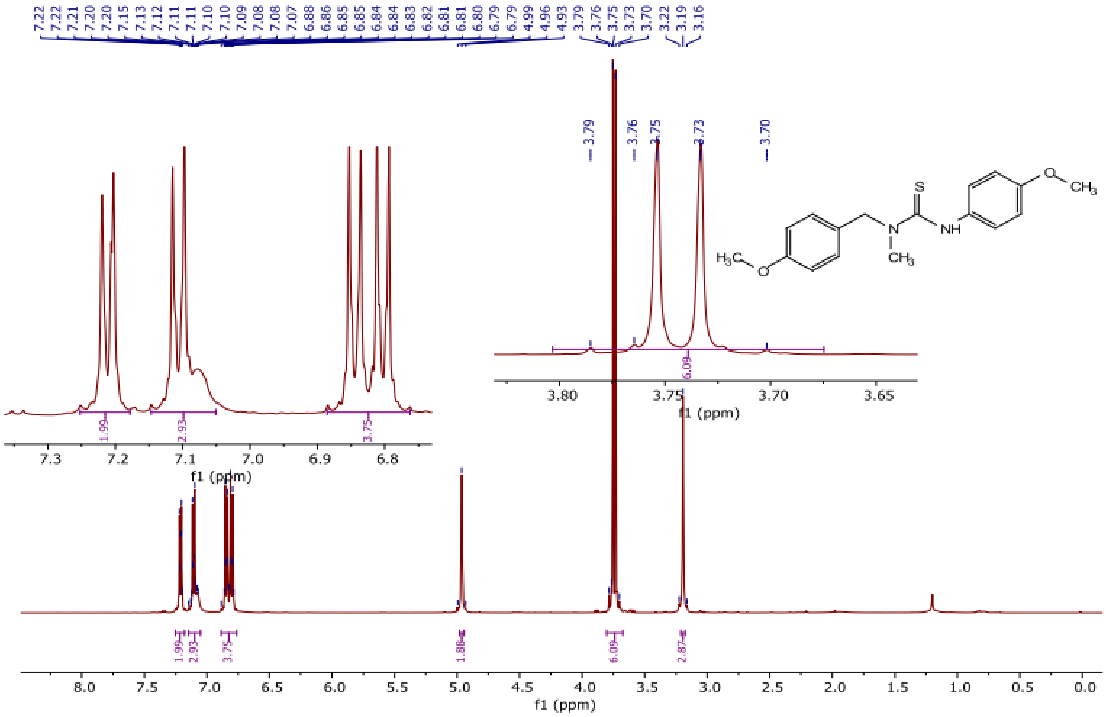
7.22 7.22 7.21 7.20 7.20 7.15 7.13 7.12 7.11 7.11 7.10 7.09 7.08 7.08 7.07 6.88 6.86 6.85 6.85 6.84 6.84 6.83 6.82 6.81 6.81 6.80 6.79 6.79 4.99 4.96 4.93 3.79 3.76 3.75 3.73 3.70 3.22 3.19 3.16
3.79
3.76
3.75
3.73
3.70
6.09
3.80 3.75 3.70 3.65
f1 (ppm)
1.99
2.93
3.75
7.3 7.2 7.1 7.0 6.9 6.8
f1 (ppm)
S
NH
N
O
CH3
H3C
O
CH3
1.99 2.93 3.75 1.88 6.09 2.87
8.0 7.5 7.0 6.5 6.0 5.5 5.0 4.5 4.0 3.5 3.0 2.5 2.0 1.5 1.0 0.5 0.0
f1 (ppm)

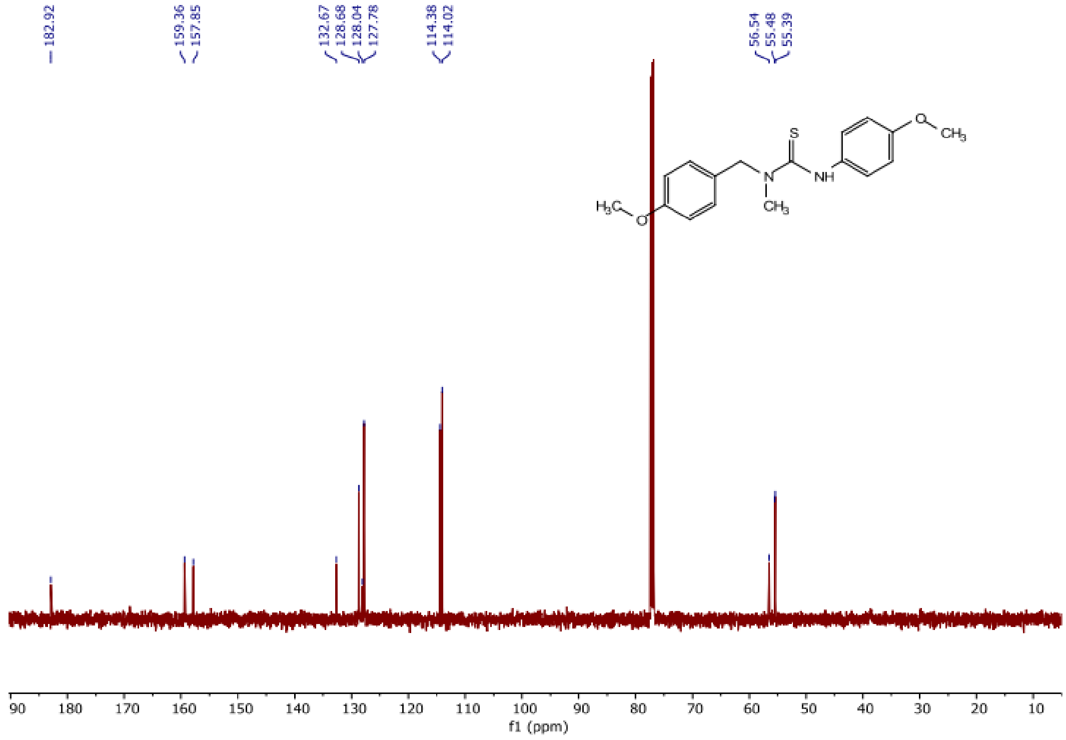
182.92
159.36 157.85
132.67 128.68 128.04 127.78
114.38 114.02
56.54 55.48 55.39
S
NH
N
O
CH3
H3C
O
CH3
90 180 170 160 150 140 130 120 110 100 90 80 70 60 50 40 30 20 10
f1 (ppm)

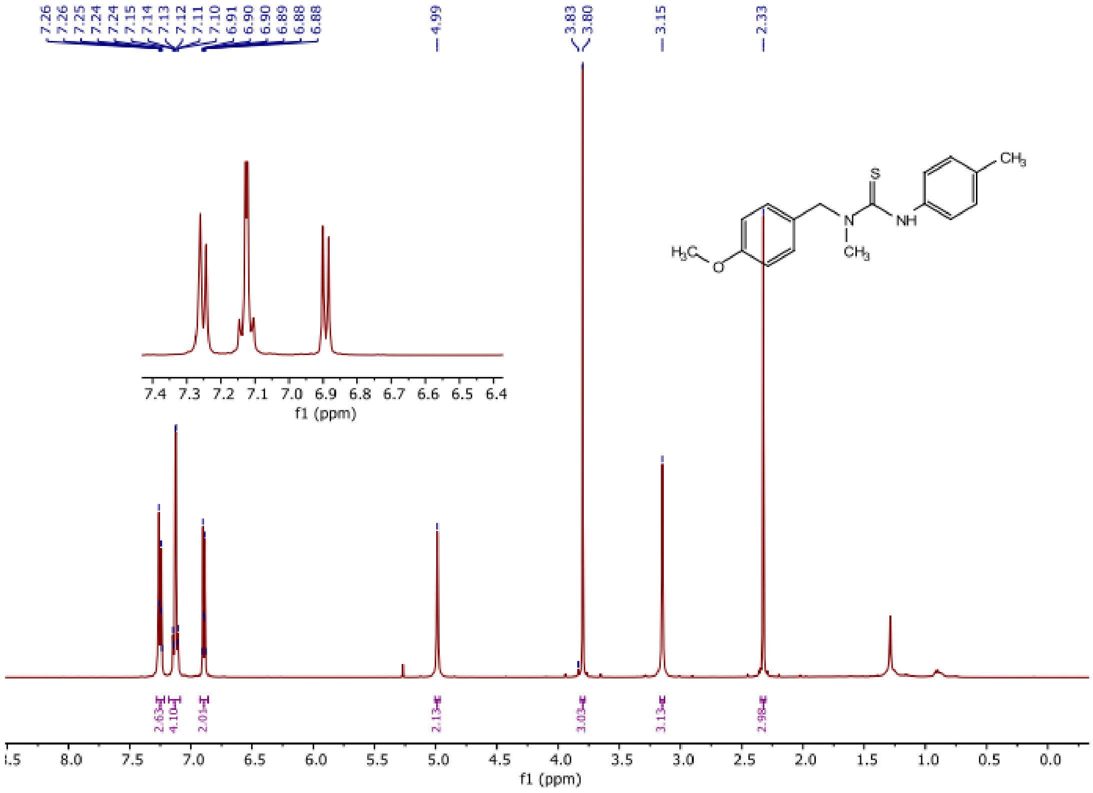
7.26 7.26 7.25 7.24 7.24 7.15 7.14 7.13 7.12 7.11 7.10 6.91 6.90 6.90 6.89 6.88 6.88
4.99
3.83 3.80
3.15
2.33
7.4 7.3 7.2 7.1 7.0 6.9 6.8 6.7 6.6 6.5 6.4
f1 (ppm)
2.63 4.10 2.01 2.13 3.03 3.13 2.98
[illegible].5 8.0 7.5 7.0 6.5 6.0 5.5 5.0 4.5 4.0 3.5 3.0 2.5 2.0 1.5 1.0 0.5 0.0
f1 (ppm)

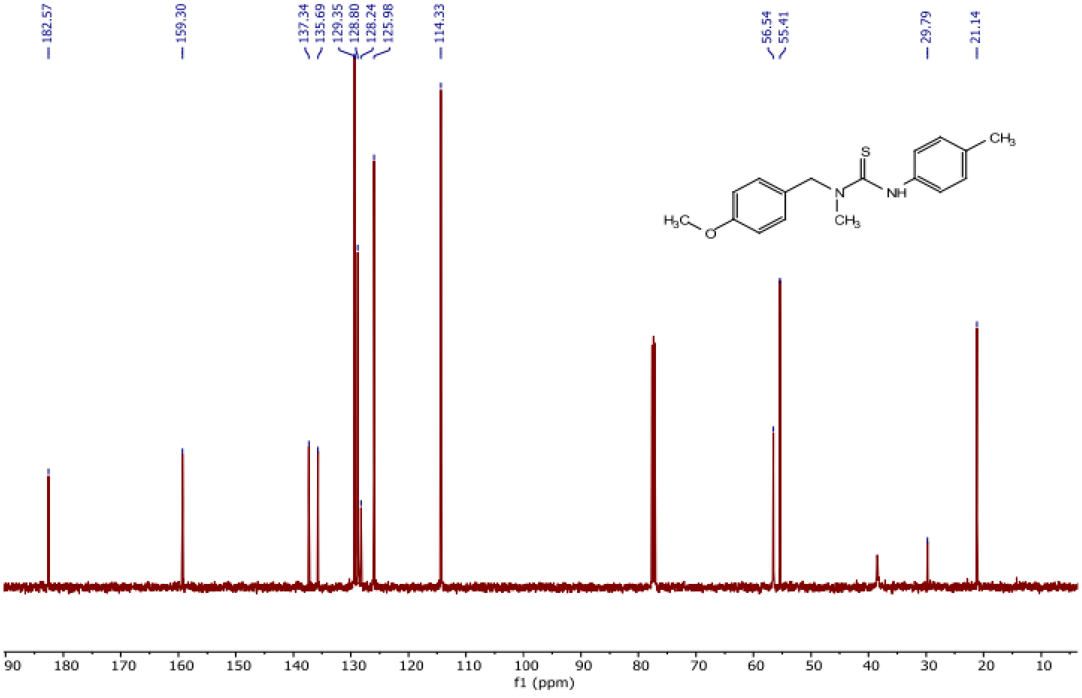
182.57
159.30
137.34 135.69 129.35 128.80 128.24 125.98
114.33
56.54 55.41
29.79
21.14
90 180 170 160 150 140 130 120 110 100 90 80 70 60 50 40 30 20 10
f1 (ppm)

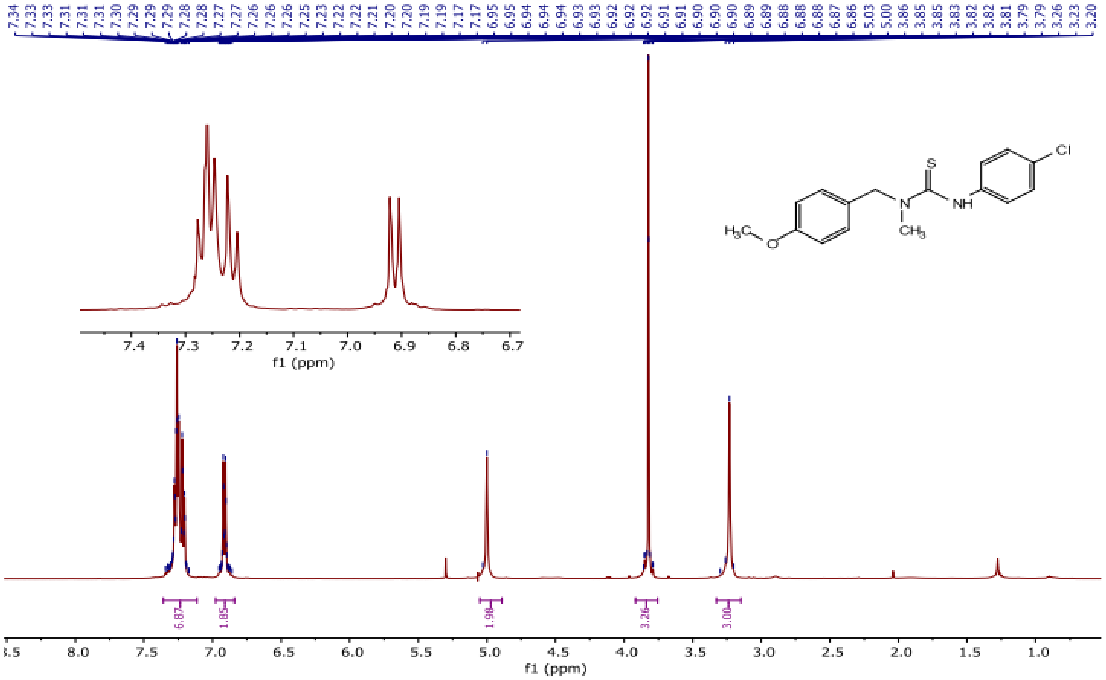
7.34 7.33 7.33 7.31 7.31 7.31 7.30 7.29 7.29 7.29 7.28 7.28 7.27 7.27 7.26 7.26 7.26 7.25 7.23 7.22 7.22 7.21 7.20 7.20 7.19 7.19 7.17 7.17 6.95 6.95 6.94 6.94 6.94 6.93 6.93 6.92 6.92 6.92 6.91 6.91 6.90 6.90 6.90 6.89 6.89 6.88 6.88 6.88 6.87 6.86 5.03 5.00 3.86 3.85 3.85 3.83 3.82 3.82 3.81 3.79 3.26 3.23 3.20
S
Cl
N
NH
$H_3C$
O
$CH_3$
6.87
1.85
1.98
3.26
3.00
f1 (ppm)
f1 (ppm)

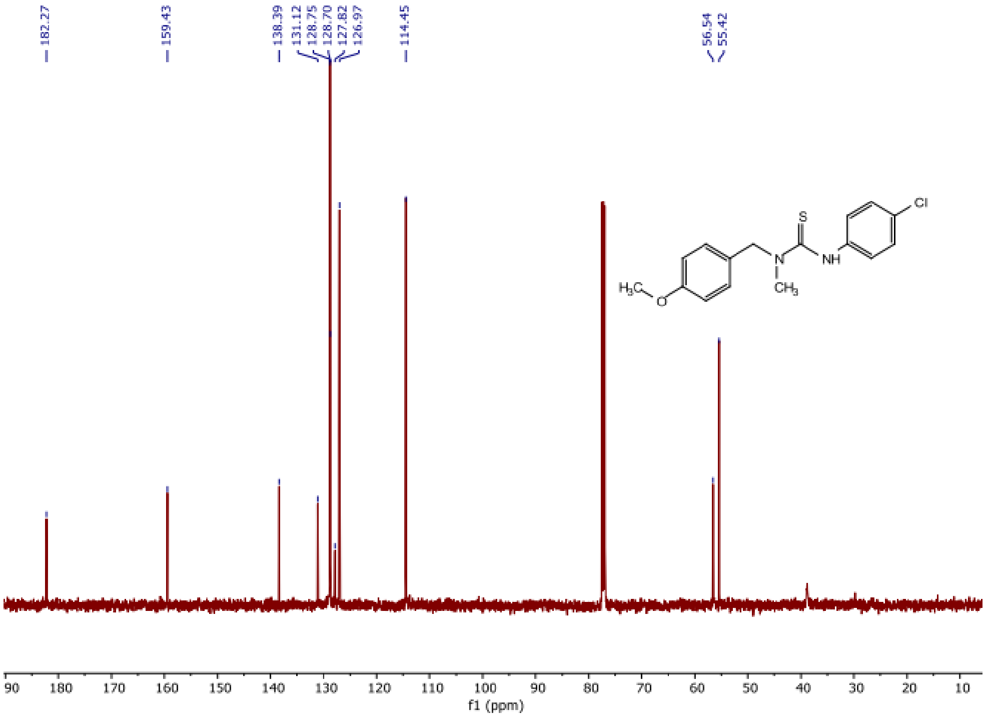
182.27
159.43
138.39 131.12 128.75 128.70 127.82 126.97
114.45
56.54 55.42
S
Cl
N
NH
$H_3C$
O
$CH_3$
f1 (ppm)

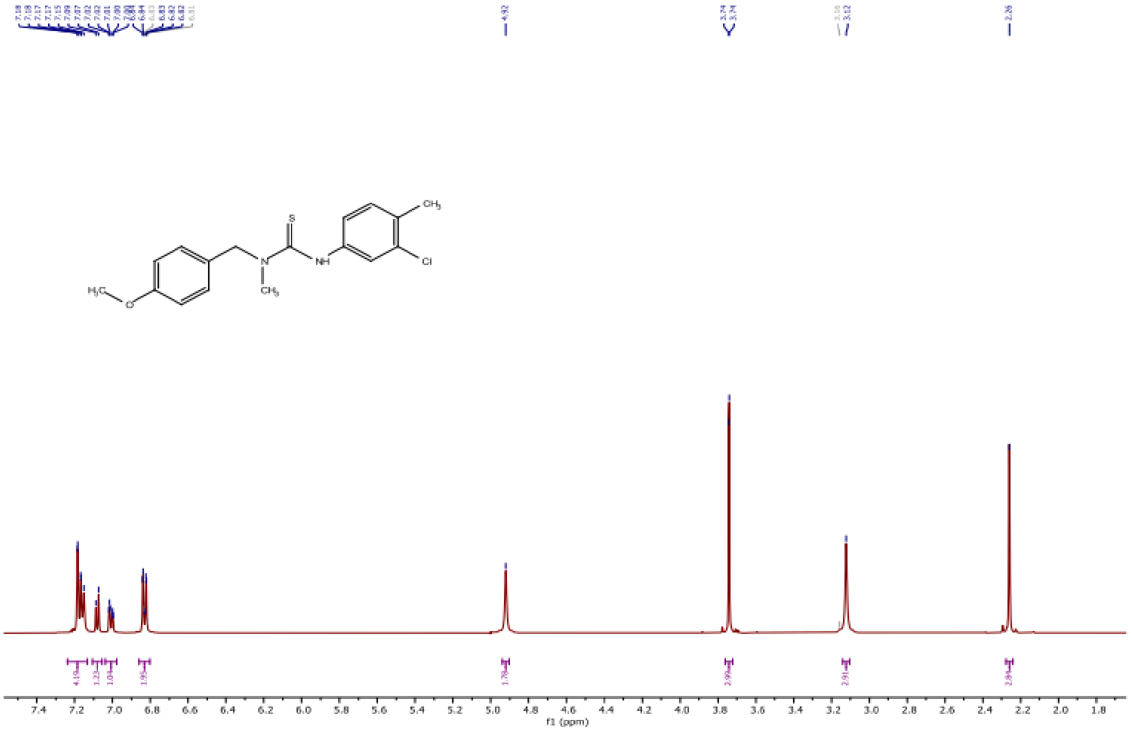
CH3
S
NH
Cl
N
CH3
H3C
O
f1 (ppm)

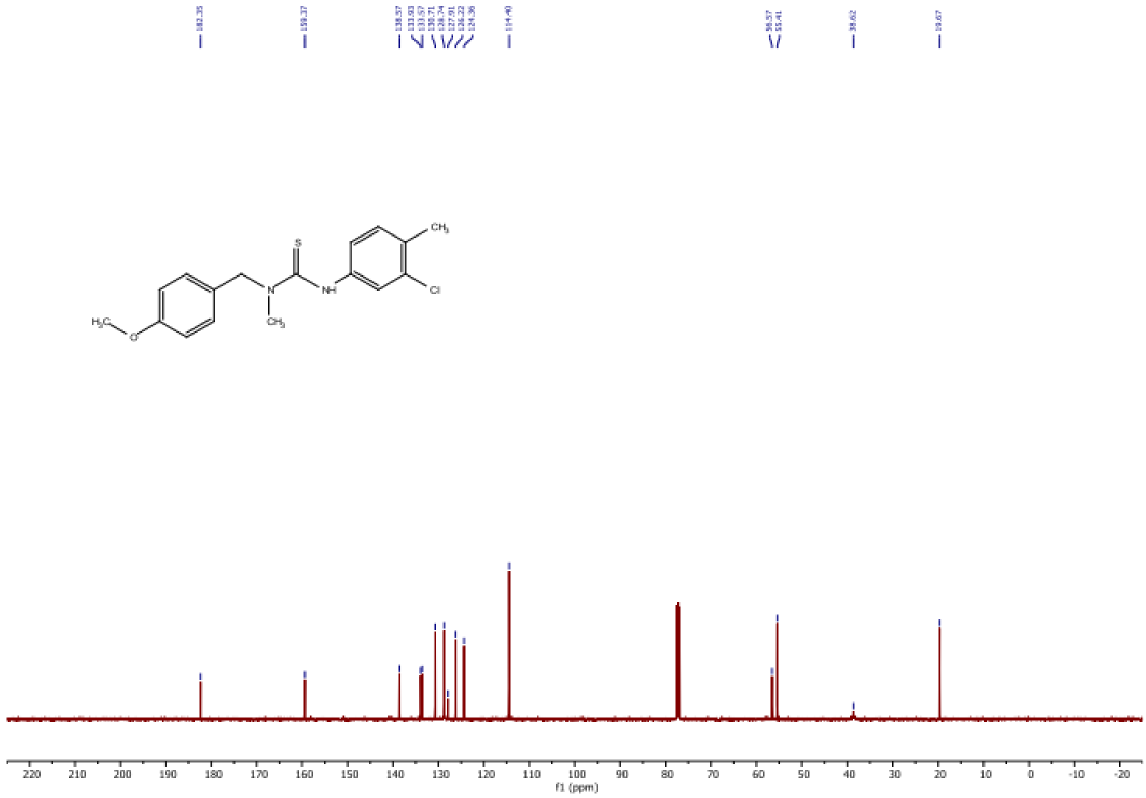
CH3
S
NH
Cl
N
CH3
H3C
O
f1 (ppm)

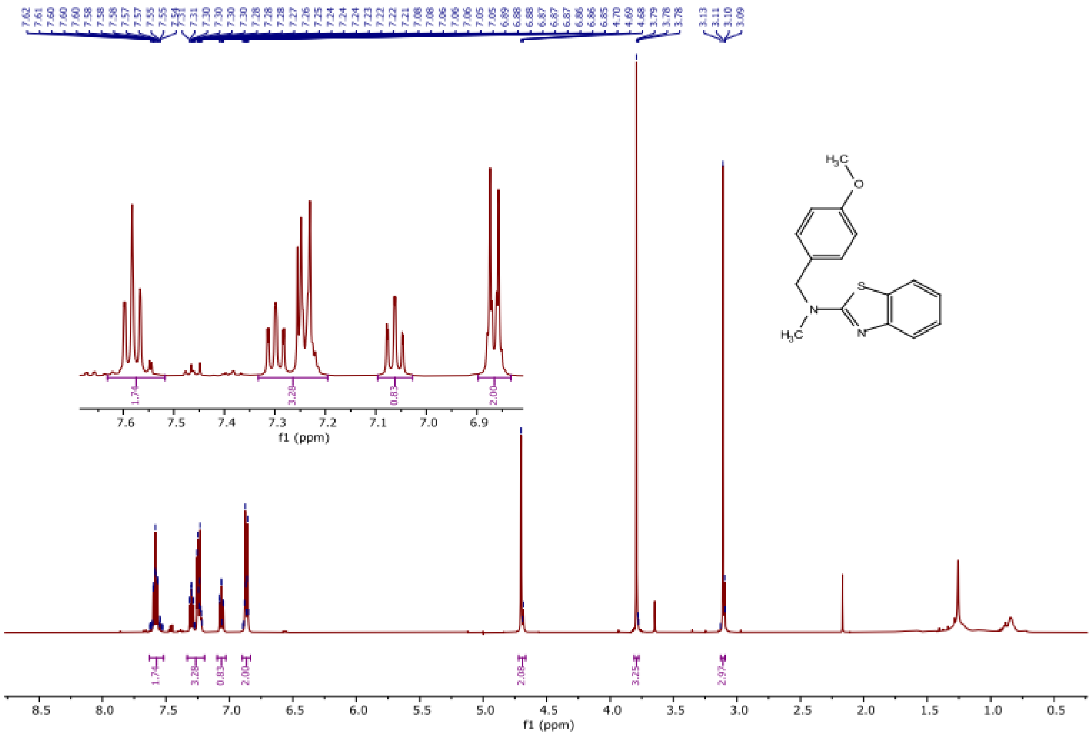
7.62 7.61 7.60 7.60 7.60 7.58 7.58 7.57 7.55 7.55 7.54 7.53 7.31 7.30 7.30 7.30 7.30 7.28 7.28 7.28 7.27 7.26 7.25 7.24 7.24 7.24 7.23 7.22 7.22 7.21 7.08 7.08 7.06 7.06 7.06 7.05 7.05 6.89 6.88 6.88 6.87 6.87 6.87 6.86 6.86 6.85 4.70 4.69 4.68 3.79 3.78 3.78 3.13 3.11 3.10 3.09
1.74
3.28
0.83
2.00
2.08
3.25
2.97
f1 (ppm)
H3C
O
S
N
N
H3C

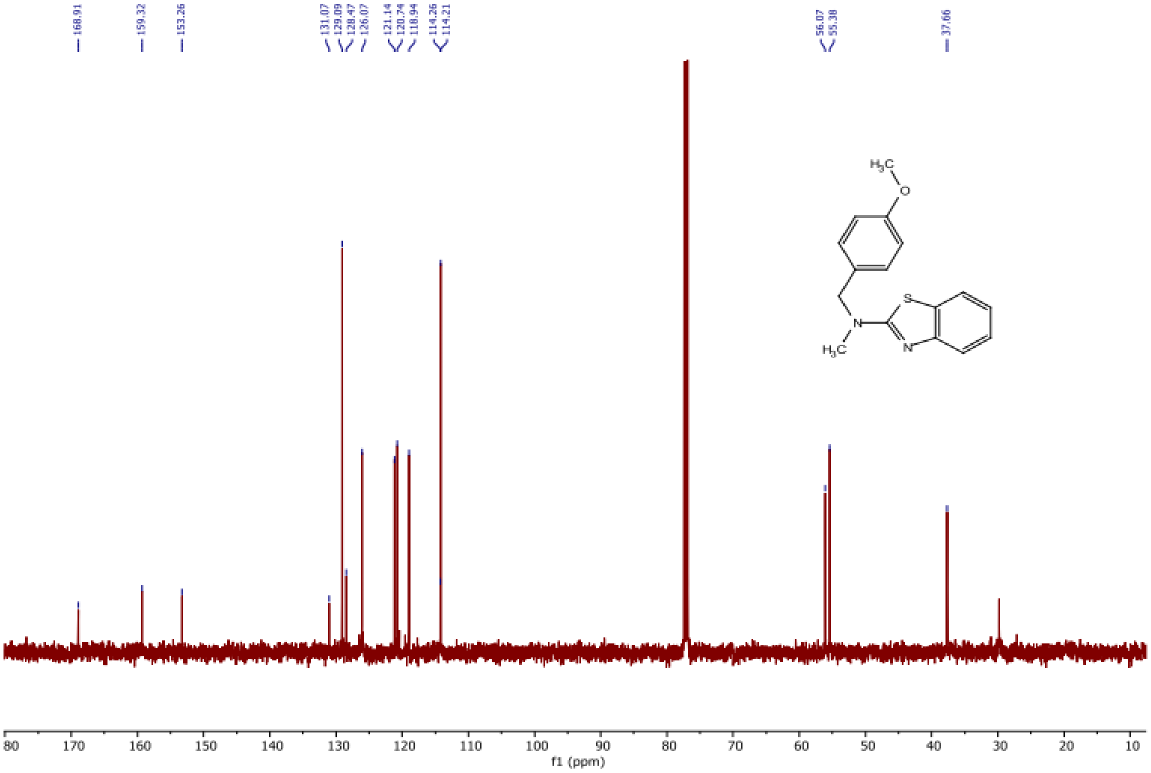
168.91
159.32
153.26
131.07 129.09 128.47 126.07
121.14 120.74 118.94
114.26 114.21
56.07 55.38
37.66
f1 (ppm)
H3C
O
S
N
N
H3C

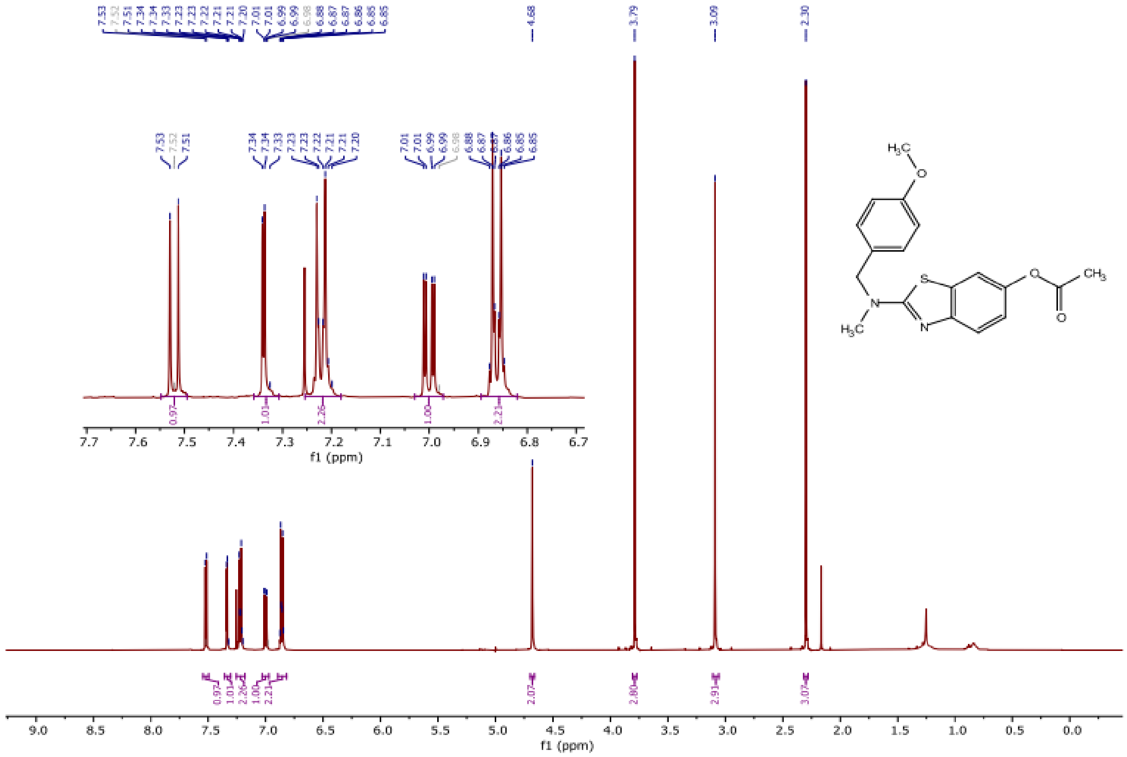
7.53
7.52
7.51
7.34
7.34
7.33
7.23
7.23
7.22
7.21
7.21
7.20
7.01
7.01
6.99
6.99
6.98
6.88
6.87
6.87
6.86
6.85
6.85
4.68
3.79
3.09
2.30
0.97
1.01
2.26
1.00
2.21
2.07
2.80
2.91
3.07
f1 (ppm)
S
N
O
CH3
H3C


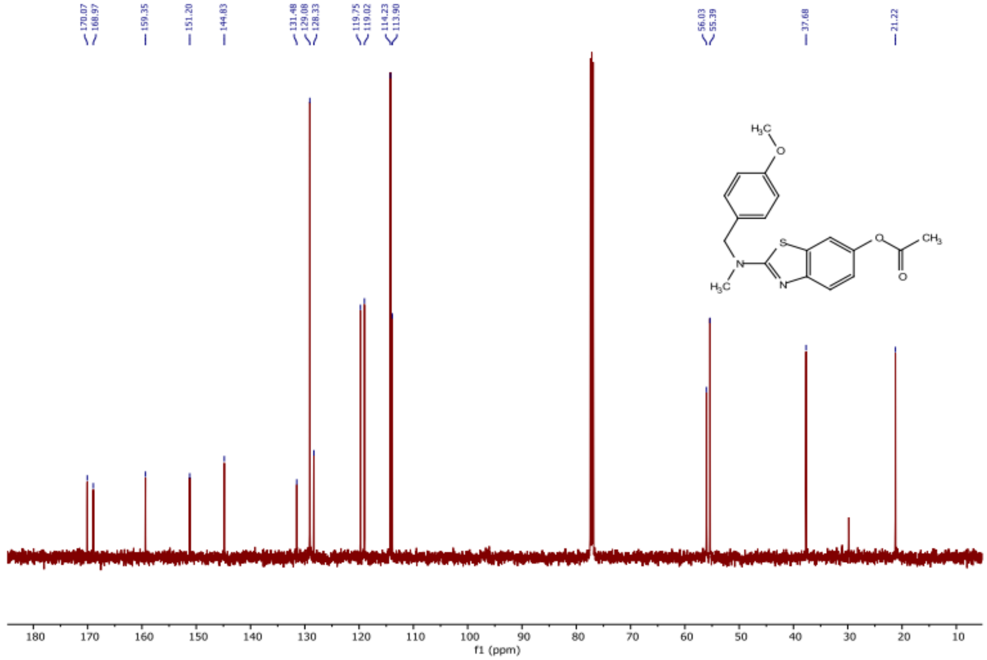
170.07
168.97
159.35
151.20
144.83
131.48
129.08
128.33
119.75
119.02
114.23
113.90
56.03
55.39
37.68
21.22
f1 (ppm)
S
N
O
CH3
H3C

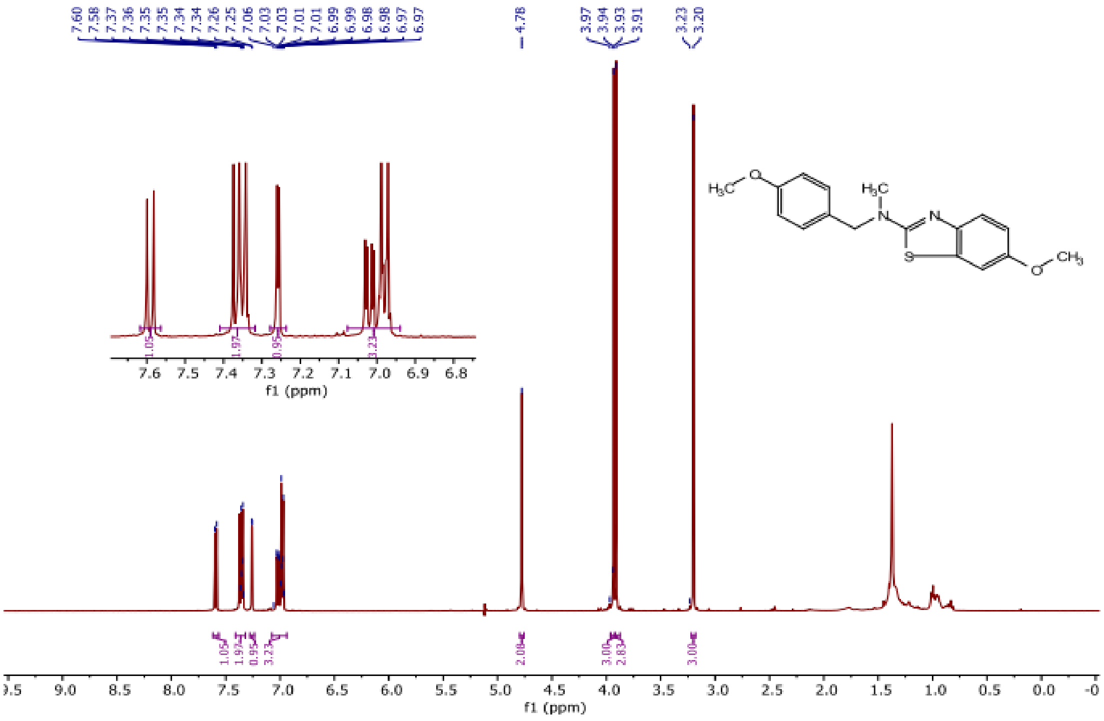

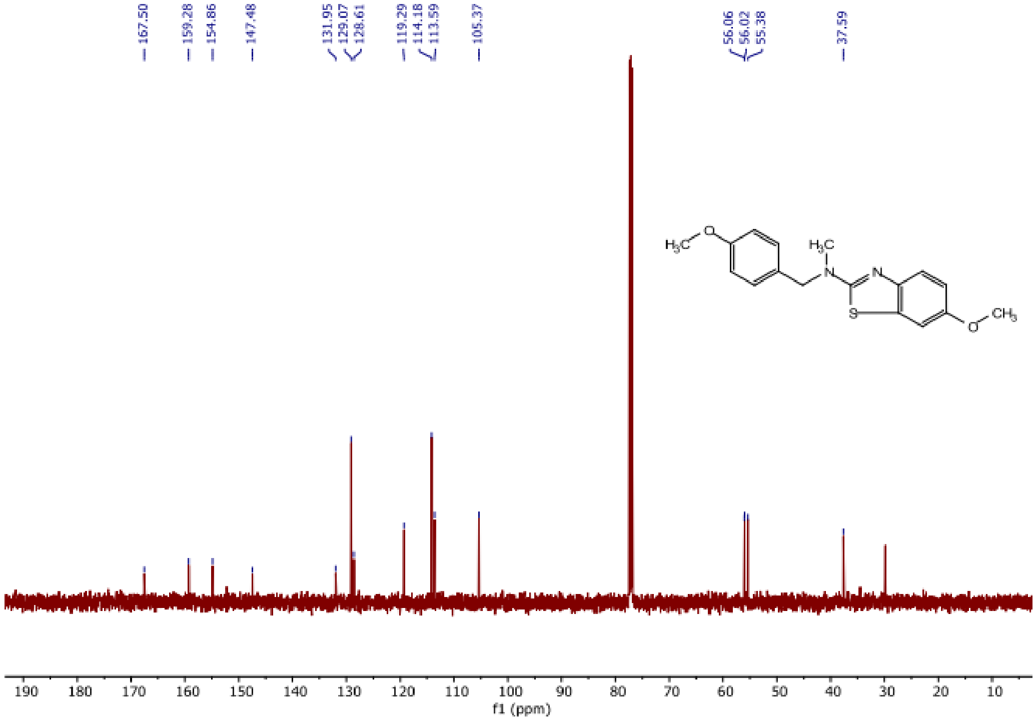

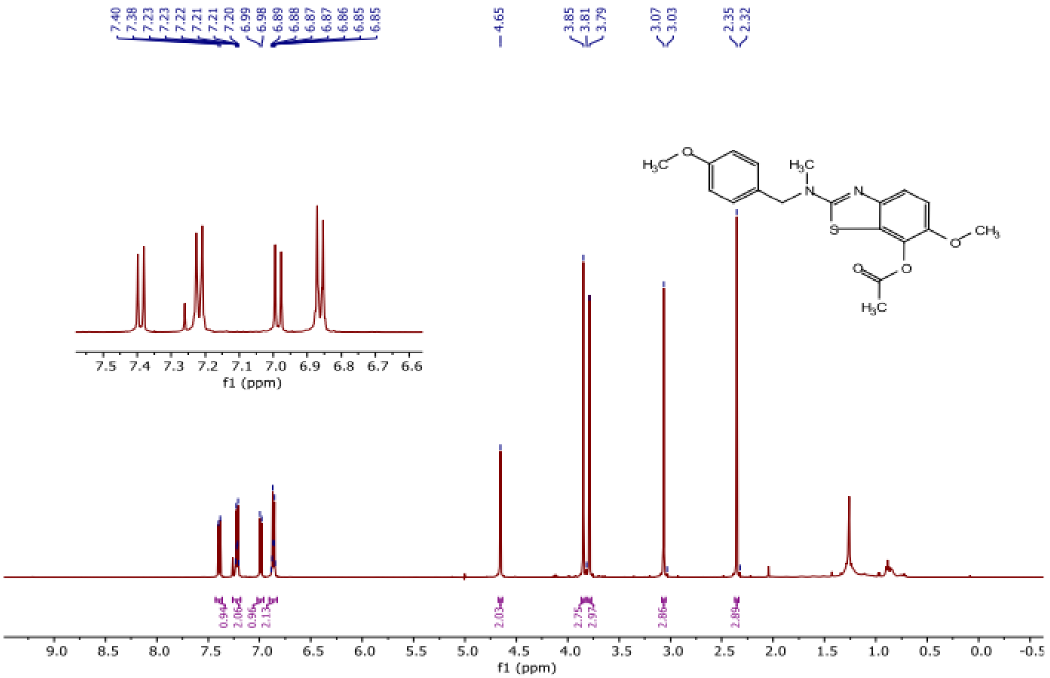

7.40
7.38
7.23
7.23
7.22
7.21
7.21
7.20
6.99
6.98
6.89
6.88
6.87
6.87
6.86
6.85
6.85
4.65
3.85
3.81
3.79
3.07
3.03
2.35
2.32
0.94
2.06
0.96
2.13
2.03
2.25
2.97
2.86
2.89
f1 (ppm)
f1 (ppm)


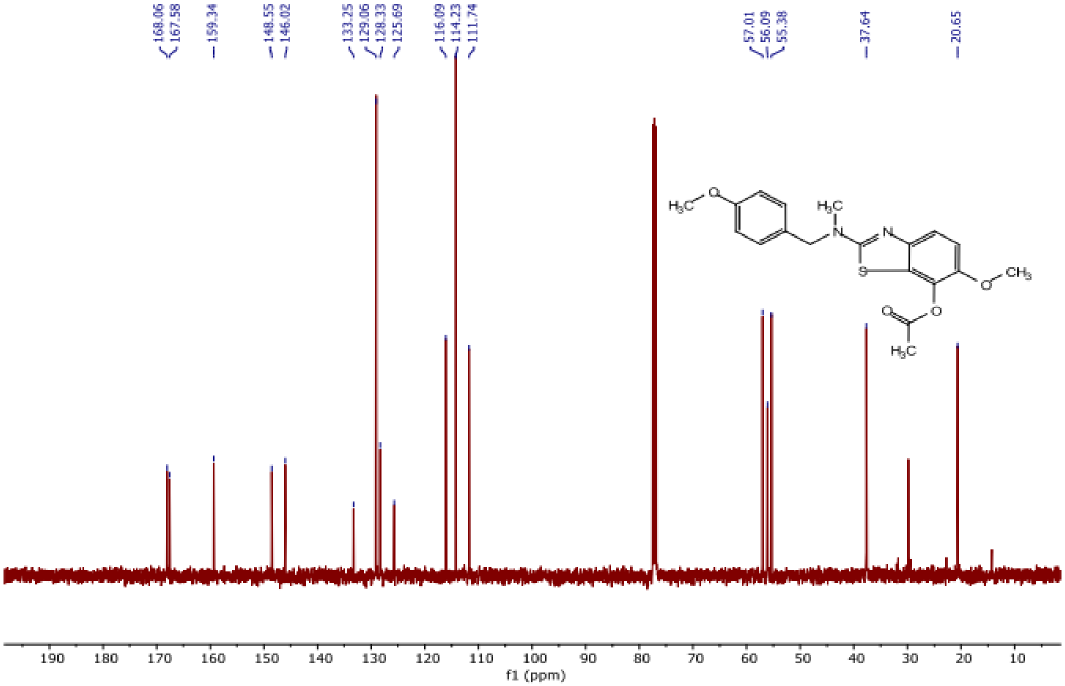

168.06
167.58
159.34
148.55
146.02
133.25
129.06
128.33
125.69
116.09
114.23
111.74
57.01
56.09
55.38
37.64
20.65
f1 (ppm)

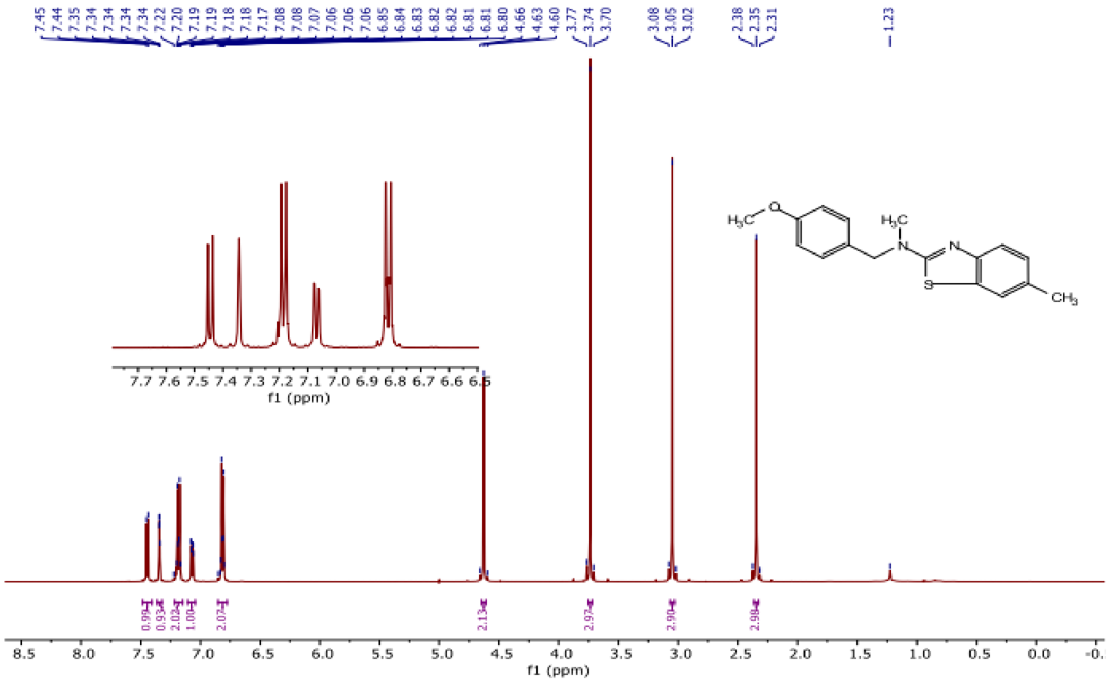
7.45
7.44
7.35
7.34
7.34
7.34
7.34
7.22
7.20
7.19
7.19
7.18
7.18
7.17
7.08
7.08
7.07
7.06
7.06
7.06
6.85
6.84
6.83
6.82
6.82
6.81
6.81
6.80
4.66
4.63
4.60
3.77
3.74
3.70
3.08
3.05
3.02
2.38
2.35
2.31
1.23
H3C
O
H3C
N
N
S
CH3
7.7 7.6 7.5 7.4 7.3 7.2 7.1 7.0 6.9 6.8 6.7 6.6 6.5
f1 (ppm)
0.99
0.93
2.02
1.00
2.07
2.13
2.97
2.90
2.98
8.5 8.0 7.5 7.0 6.5 6.0 5.5 5.0 4.5 4.0 3.5 3.0 2.5 2.0 1.5 1.0 0.5 0.0 -0.
f1 (ppm)

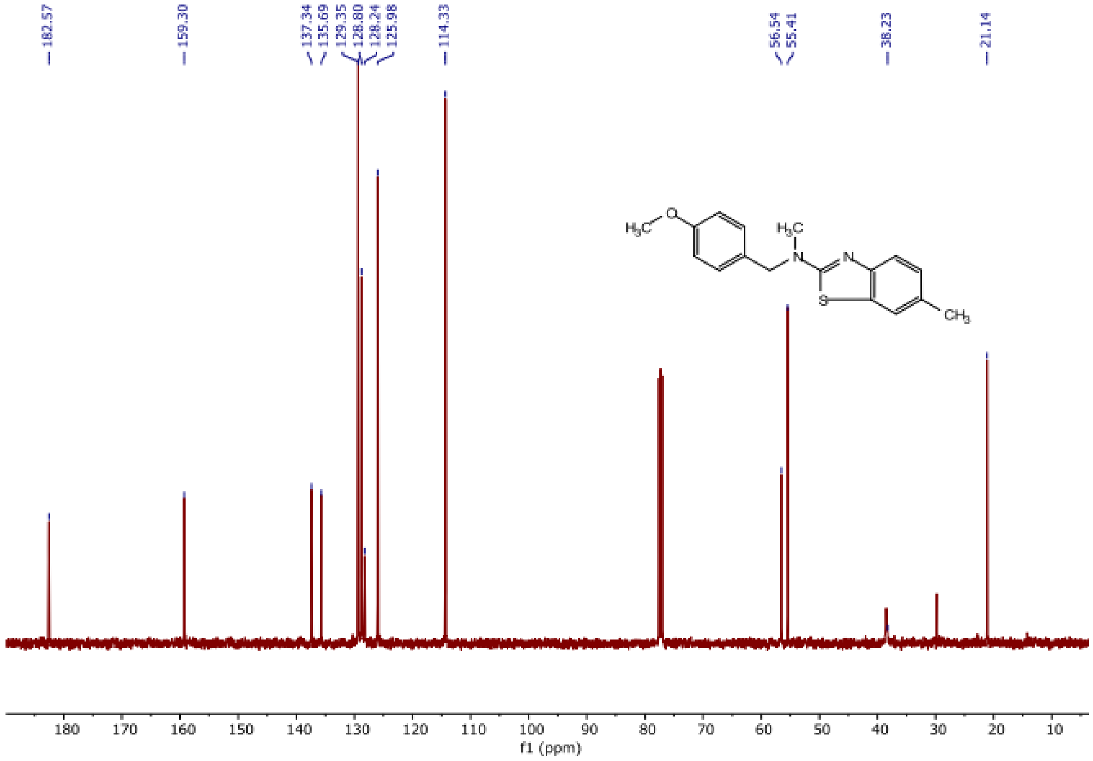
182.57
159.30
137.34
135.69
129.35
128.80
128.24
125.98
114.33
56.54
55.41
38.23
21.14
H3C
O
H3C
N
N
S
CH3
180 170 160 150 140 130 120 110 100 90 80 70 60 50 40 30 20 10
f1 (ppm)

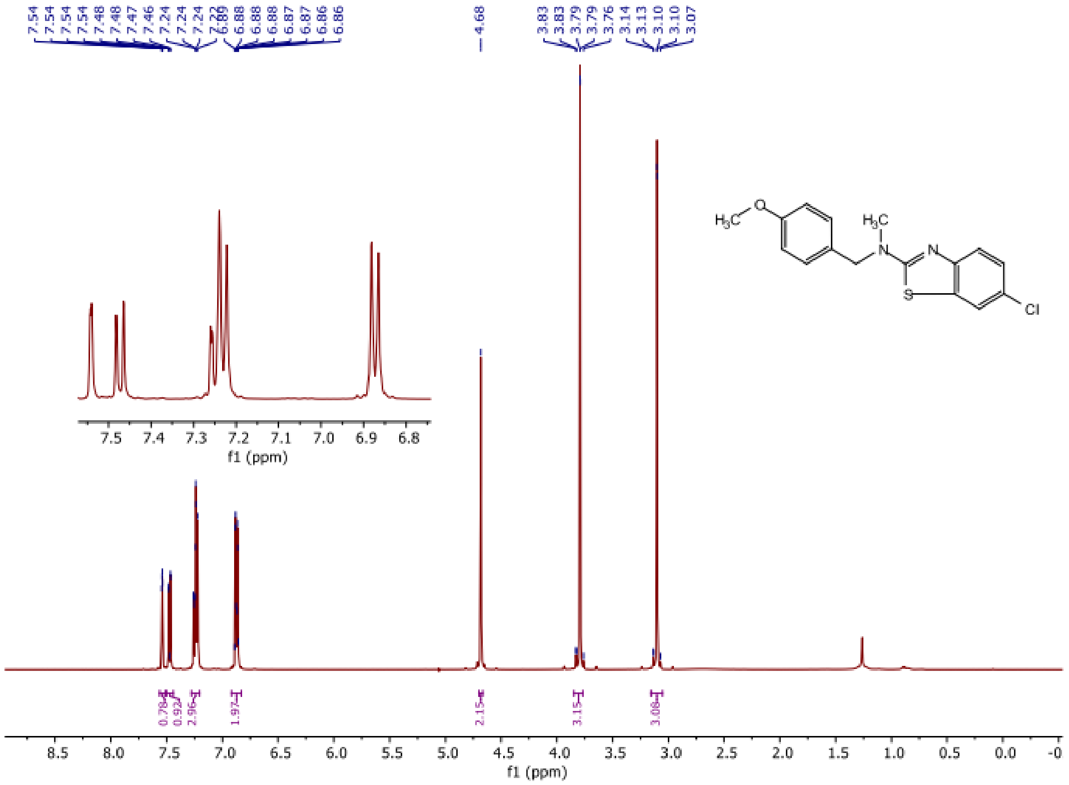
7.54 7.54 7.54 7.54 7.48 7.48 7.47 7.46 7.24 7.24 7.24 7.22 6.89 6.88 6.88 6.88 6.87 6.86 6.86
4.68
3.83 3.83 3.79 3.79 3.76 3.14 3.13 3.10 3.10 3.07
0.79 0.92 2.96 1.97 2.15 3.15 3.08
f1 (ppm)
H3C O H3C N N S Cl

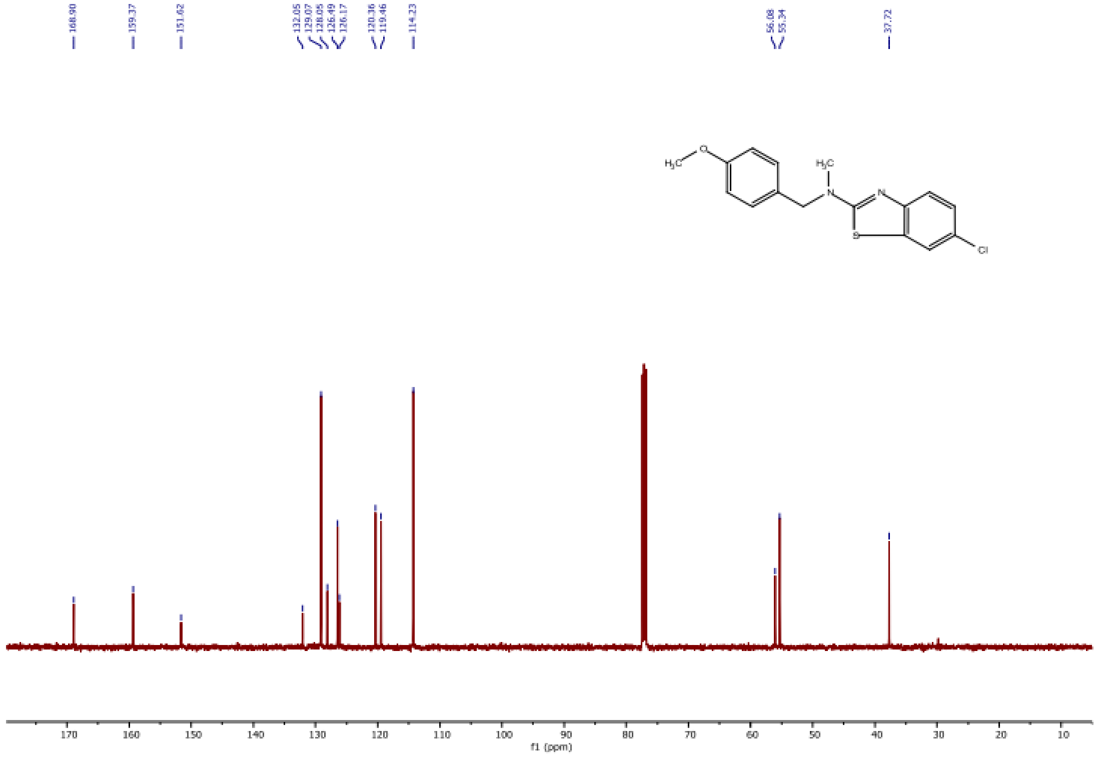
168.90
159.37
151.62
132.05 129.07 128.05 126.49 126.17
120.36 119.46
114.23
56.08 55.34
37.72
f1 (ppm)
H3C O H3C N N S Cl

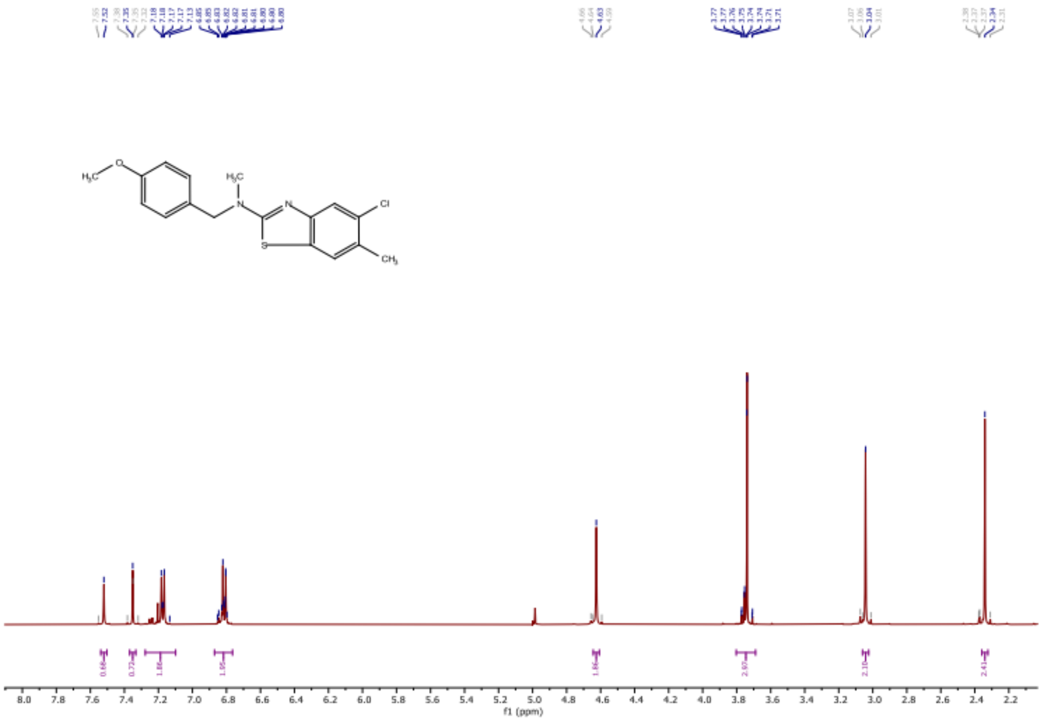

H3C
O
H3C
N
N
S
Cl
CH3
f1 (ppm)


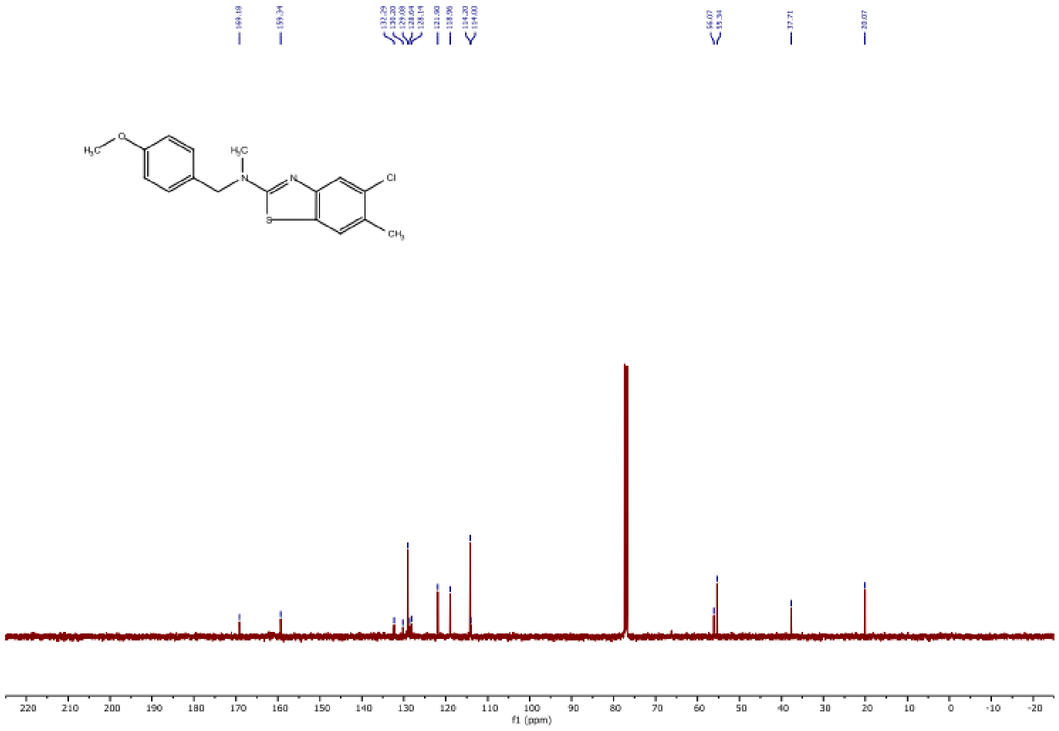

H3C
O
H3C
N
N
S
Cl
CH3
f1 (ppm)
